# Accuracy of occurrence and abundance estimates from insect metabarcoding

**DOI:** 10.64898/2026.02.20.707016

**Authors:** Ela Iwaszkiewicz-Eggebrecht, Emma Granqvist, Karol H. Nowak, Catalina Valdivia, Mateusz Buczek, Amrita Srivathsan, Emily Hartop, Andreia Miraldo, Tomas Roslin, Ayco J. M. Tack, Piotr Łukasik, Rudolf Meier, Fredrik Ronquist

**Author notes:** <u>Correspondence author</u>.

## Abstract

1. DNA metabarcoding—high-throughput sequencing of barcode regions from bulk samples—has become a key tool for insect biodiversity assessment. Yet, how methodological choices affect the accuracy of metabarcoding data remains insufficiently explored. In this paper, we ask: (1) How does the lysis method (non-destructive lysis vs. destructive homogenization) affect community recovery? (2) How comprehensively does metabarcoding capture species richness? (3) To what extent can spike-ins improve abundance estimates? (4) How accurately can species abundances be estimated?
2. We evaluated the accuracy of insect metabarcoding using 4,749 bulk samples from a large-scale biodiversity survey subjected to mild lysis. Of these samples, 856 were also homogenized, allowing a systematic comparison of the effect of alternative treatments. To potentially improve abundance estimates, we added six biological spike-ins (i.e., foreign insects) to all samples, and two synthetic spike-ins (artificial DNA fragments) to the homogenization treatment. In addition, we established the contents of 15 samples by individually barcoding all specimens, enabling direct assessment of occurrence and abundance estimates.
3. Our results revealed consistent differences between destructive and non-destructive treatments. While both methods reliably detected the majority of species, small and soft-bodied taxa were more often recovered after mild lysis than after homogenization, while the reverse was true for heavily sclerotized, hairy, and large taxa. Using biological spike-ins for calibration reduced the variance in read numbers per specimen considerably, especially in homogenized samples, while synthetic spike-ins were less effective. In a Bayesian analysis, where species data were matched to the best-fitting spike-in calibration curve, accurate abundance estimates (+/-1 individual) were obtained for 72.9% of species occurrences.
4. Our results show that it is possible to obtain reasonably accurate abundance estimates from metabarcoding data, and that mild lysis and homogenization result in different taxon-specific biases in terms of occurrence data, with neither method outperforming the other. Accuracy is improved by homogenization rather than mild lysis of samples, and by the use of biological rather than synthetic spike-ins. Together, these findings provide a major step towards robust, quantitative biodiversity monitoring using DNA-metabarcoding.

## Introduction

Insects play a pivotal role in the world’s ecosystems, influencing diverse ecological processes, securing food production and serving as crucial indicators of environmental health (Roslin, 2024; Scudder, 2017). As our planet experiences escalating anthropogenic pressures and climate change, insect biodiversity is increasingly threatened, rendering the need for robust and efficient monitoring tools ever more urgent (Klink et al., 2020; Wagner et al., 2021). However, traditional methods of insect monitoring rely on time-consuming manual collection and morphological identification. Due to the scarcity of taxonomic expertise and the sheer volume of specimens collected, such methods will struggle to capture the full spectrum of biodiversity (Montgomery et al., 2021; Ronquist et al., 2020).

In this context, DNA metabarcoding - a high-throughput technique for sequencing marker region amplicons for multi-species samples - has emerged as a transformative approach, offering rapid and cost-effective biodiversity assessments (Blackman et al., 2019; Hajibabaei et al., 2019). By amplifying and sequencing short, target DNA regions, metabarcoding enables the simultaneous identification of multiple species from complex bulk samples (Braukmann et al., 2019). Originally developed for microbial communities, metabarcoding has found increasing applications in insect monitoring, providing valuable insights into species composition and distribution (Åström et al., 2025; Buchner et al., 2024). Its broad implementation across national-scale projects using standardized trapping networks could greatly accelerate insect biodiversity discovery (Iwaszkiewicz-Eggebrecht et al., 2025).

Despite the obvious advantages of metabarcoding, its limitations remain underexplored. Chief among these concerns is the method’s ability to accurately recover species occurrence and abundance in complex bulk samples. Technical aspects – such as preservation of the sample, DNA extraction variability, primer bias, amplification efficiency, and uneven reference library coverage – can distort diversity estimates (Elbrecht & Leese, 2015; Alberdi et al., 2018; Elbrecht et al., 2019). Biological aspects – i.e., variation in DNA yield due to body size, degree of sclerotization, or mitochondrial copy number – may further cause some taxa to be overrepresented in sequence yields while others remain undetected (Iwaszkiewicz-Eggebrecht et al., 2023).

One critical source of methodological variation during metabarcoding lies in the choice of DNA extraction protocol. Two widely used approaches – mild lysis and homogenization – differ fundamentally in their implications for downstream analyses. In the mild lysis protocol, insect specimens are incubated in a lysis buffer without undergoing mechanical disruption, allowing DNA to leak out of their bodies (Batovska et al., 2021; Carew et al., 2018; Nielsen et al., 2019). This non-destructive method preserves the integrity of specimens for subsequent morphological or genetic work and is particularly valuable when dealing with previously uncharacterized taxa. In contrast, the homogenization approach involves physically grinding specimens into a homogenate mixture or “insect soup”, resulting in higher DNA yields from larger and sclerotized insects (Yu et al., 2012; Zizka et al., 2022). It does, however, come at the cost of destroying specimens, preventing their subsequent use for taxonomic or molecular work, and complicating the verification of the accuracy of metabarcoding results. It also comes with the risk of losing the capability to detect small species that contribute very little DNA to the “insect soup” (Elzbieta Iwaszkiewicz-Eggebrecht, Granqvist, et al., 2023). These contrasting trade-offs have fueled ongoing debate, but the performance of alternative approaches when applied to real-world samples remains insufficiently tested.

Studies comparing these protocols have often relied on mock communities – i.e., artificial mixtures of known species – as benchmarks (Iwaszkiewicz-Eggebrecht, Granqvist, et al., 2023; Luo et al., 2023). While these controlled systems have proven valuable for method development, they fail to replicate the biological and taxonomic complexity of genuine insect bulk samples. Other studies have attempted to benchmark mild lysis and homogenization using real field samples (Kirse et al., 2023), but these efforts often lack data on the species composition of the specimens in the samples, making it difficult to assess the accuracy of detection or abundance estimates. Consequently, the true performance and limitations of metabarcoding in natural insect communities – where hundreds or thousands of species co-occur – remain poorly understood. This *status quo* highlights the need for comprehensive benchmarking against real insect samples of known composition to evaluate detection accuracy and inform best practices for future biodiversity monitoring.

Beyond information on species presence-absence, quantitative data on species abundance and biomass are central to biomonitoring efforts and form part of the Essential Biodiversity Variables framework used to track ecological change (Pereira et al., 2013). Abundance and biomass together shape species interactions, community dynamics, and ecosystem functioning and are essential for predicting how insect communities respond to environmental change (Uhler et al., 2021). Yet for insects, it is notoriously difficult to obtain precise quantitative information. Manual sorting, counting and weighing thousands of individuals from bulk samples is prohibitively time-consuming and expensive, requires extensive taxonomic expertise, which is itself scarce, and is ultimately infeasible for large-scale monitoring programs (Karlsson et al., 2020) - i.e., for the very programs most needed to track biodiversity change. Automated computer vision-based methods promise to change the situation, but remain to be effectively implemented at a large scale (Høye et al., 2021).

While not originally designed for quantitative inference, metabarcoding offers a potential pathway toward abundance and biomass estimation through the use of read numbers. This is clearly a challenging problem, as it is well known that read counts are affected by specimen variation, as well as by species- and sample-specific factors (Iwaszkiewicz-Eggebrecht, Granqvist, et al., 2023; Martoni et al., 2022). While variation among specimens is difficult to address, species- and sample-specific factors might be controlled for by using spike-ins (i.e., standards added at known quantities) for calibration (Ji et al., 2020; Luo et al., 2023; Sickel et al., 2023). Spike-ins can be implemented in different forms, each with distinct advantages. Biological spike-ins - i.e., individuals of species absent from the focal region - are added to samples early in the workflow, providing a reference for how DNA recovery and amplification vary across samples. In addition, if several spike-ins of several species with different characteristics are used, they can also capture how DNA recovery varies across species, depending on factors such as different body size and degree of sclerotization (Iwaszkiewicz-Eggebrecht et al., 2023). Synthetic spike-ins, by contrast, are short artificial DNA sequences added in precise quantities before or after DNA extraction from the bulk sample (Harrison et al., 2021; Tourlousse et al., 2017). Because synthetic spike-ins are unambiguously recognizable and avoid primer bias by design, they can capture variation across samples in the PCR amplification and sequencing steps very precisely. Thus, these strategies offer complementary ways to assess and reduce variation in metabarcoding outputs (Iwaszkiewicz-Eggebrecht et al., 2024). Nonetheless, how their efficiencies compare and what accuracy one might expect from the calibrated abundance estimates is still poorly known .

In this study, we combine large-scale data from field-collected samples with clearcut benchmarks, in which the specimen content of individual samples is established. By this approach, we evaluate the accuracy in occurrence and abundance estimates obtained using different metabarcoding protocols. Using 856 insect samples from Sweden, we compare destructive (homogenization) and non-destructive (mild lysis) treatments. To establish a benchmark for accuracy, we combine metabarcoding with individual-based DNA barcoding of 15 Malaise trap samples, providing comparative data for species occurrences and abundances. We further assess the potential of both biological and synthetic spike-ins in improving abundance estimation using a broader set of 4,749 samples. Finally, we develop and apply a hierarchical Bayesian model that incorporates spike-in data to explore the accuracy one might expect in estimating species abundances (specimen numbers) in Malaise trap samples.

Together, these analyses allow us to rigorously test how methodological choices affect the accuracy of occurrence and abundance estimates in insect metabarcoding, bringing us closer towards integrating high-throughput molecular data into robust, quantitative biodiversity monitoring.

## Materials and Methods

### Sample collection and metabarcoding

As the material for the study, we used 4,749 weekly Malaise trap catches collected in Sweden in 2019 as part of the *Insect Biome Atlas* project (IBA, Miraldo et al., 2025). All samples were processed using non-destructive mild lysis following the FAVIS protocol (Elzbieta Iwaszkiewicz-Eggebrecht, Łukasik, et al., 2023). For a subset of samples (n=856; Fig. 1A), the insects recovered from mild lysis were subsequently homogenized together with the remaining, sample-specific lysate, as described in the protocol of Persson et al. (2024). Following the FAVIS protocol (Iwaszkiewicz-Eggebrecht, Łukasik, et al., 2023), six biological spike-in species were added to each sample before mild lysis: three species of *Drosophila* (*D. serrata, D. jambulina, D. bicornuta), Gryllodes sigillatus, Gryllus bimaculatus* and *Shelfordella lateralis* (for specimen numbers see Fig. 1B). Following homogenization and before DNA purification, we added two plasmids as synthetic spike-ins, each in an estimated 5M copies. The two plasmids carried alternative synthetic targets (*tp53-synth* and *Callio-synth*; Iwaszkiewicz-Eggebrecht et al., 2024) designed to differ from any known barcode sequence but to amplify well in metabarcoding studies. All details regarding the specific sequence and production of plasmids are described by Iwaszkiewicz-Eggebrecht, Buczek, et al. (2023). Negative controls were added at different stages of processing (20 buffer blanks, 13 DNA extraction negatives, 11 PCR negatives). In addition, we included ten positive controls at the DNA extraction step.

**Fig. 1.**
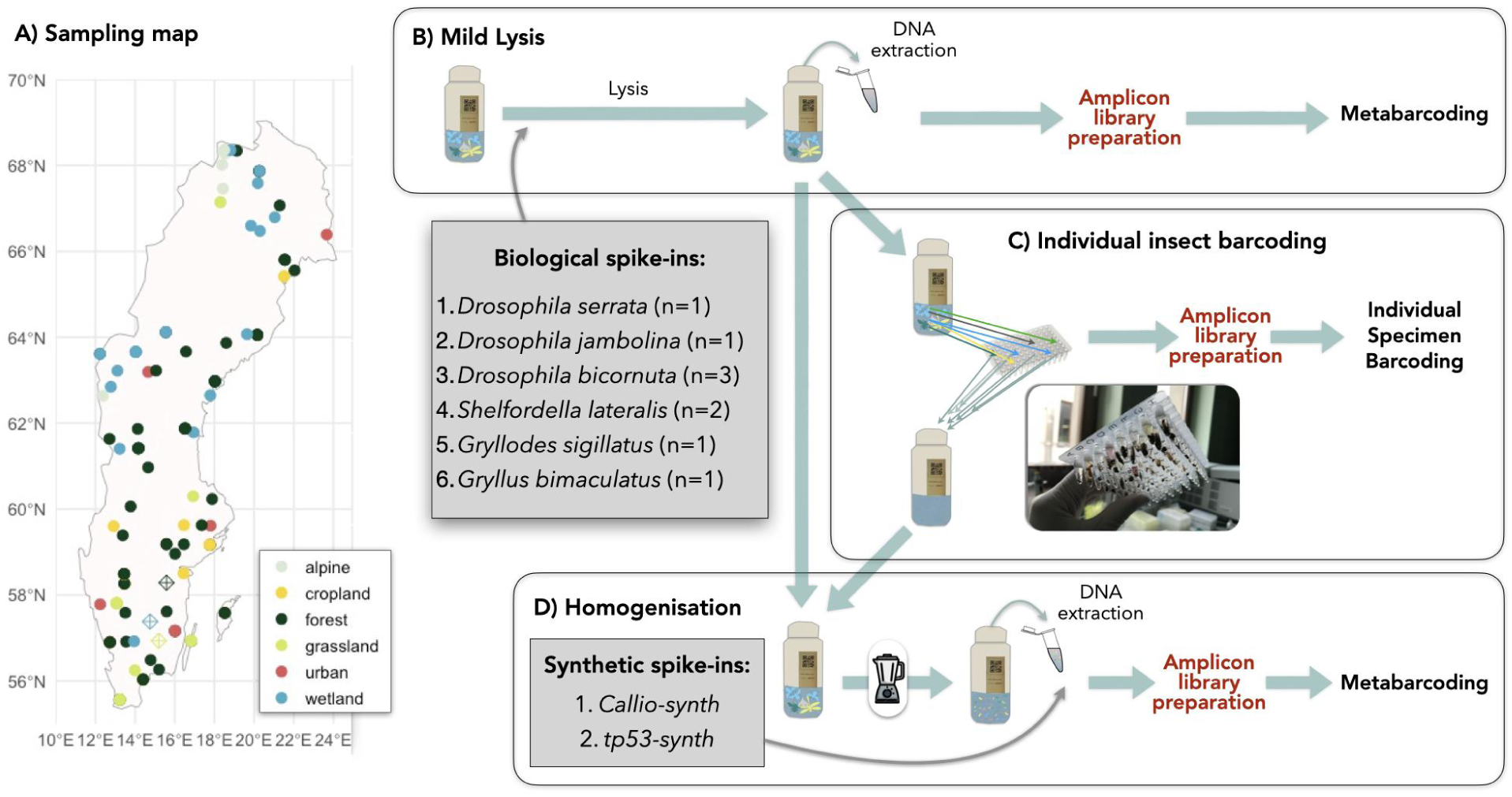
Schematic overview of the sampling locations and sample processing. (A) Map of Sweden showing locations of samples that underwent both mild lysis and homogenization (n=856). Colors indicate different habitat types. Diamonds mark the locations of the 15 samples that were additionally used for individual barcoding. (B–D) Schematic overview of sample processing. All samples (n=4,749) underwent mild lysis (B) and a subset (n=856) was subsequently homogenized (D), followed by DNA extraction and metabarcoding. Biological spike-ins were added to each lysate sample prior to mild lysis, and homogenates had additional synthetic spike-ins added before DNA extraction. A subset of 15 samples was additionally sorted to the individual level and processed for DNA barcoding (C), providing a reference dataset for validating metabarcoding results.

DNA was purified from a 225 µL aliquot of each lysate and homogenate. We then prepared amplicon libraries targeting a 418-bp long fragment of the cytochrome *c* oxidase I (COI) gene with BF3-BR2 primers (Elbrecht et al., 2019). All libraries were sequenced on an Illumina NovaSeq 6000, with SP 2×250-bp flow cells. Sequencing was performed at SciLifeLab in Stockholm (National Genomics Infrastructure Sweden) to the target coverage of 1M read pairs per library. A detailed description of molecular work can be found in Iwaszkiewicz-Eggebrecht, Buczek, et al. (2023).

Metabarcoding sequencing results were processed following the methods described by Sundh et al. (2025). In short, removal of adapters, primer trimming, and length filtering were carried out with *Cutadapt v3.1* (Martin, 2011). Reads were retained if 403–418 nt long (in 3-nt steps) and free of in-frame stop codons. Denoising was performed with the *nf-core/ampliseq Nextflow pipeline v2.4.0* (Straub et al., 2020), which applies *DADA2* (Callahan et al., 2016) to infer amplicon sequence variants (ASVs). Taxonomic annotation and further filtering were conducted with the HAPP pipeline (Sundh et al., 2025). This included taxonomic classification, chimera removal, OTU clustering, and the removal of NUMTs and low-abundance noise. Specifically, taxonomy was assigned using *SINTAX* (Edgar, 2016) implemented in *VSEARCH* (Rognes et al., 2016), against a custom COI reference database, with 80% assignment confidence set as the cutoff. The custom database was built from all BOLD sequences associated with BINs available in December 2022 (Sundh, 2023). Sequences were cleaned of gaps and ambiguous characters, and clustered at 100% identity per BIN using the *coidb* package (Sundh, 2024).

To evaluate how the processing method influenced taxonomic representation, we compared cluster counts between mild lysis and homogenization within each sample. For clusters detected in both treatments, we calculated the ratio of read counts between lysates and homogenates, applying a normalization step to account for differences in sequencing depth. Clusters were then grouped by family, and the distribution of read ratios was examined for well-represented families (Fig. 2A). Clusters detected exclusively in one treatment were likewise summarized by family (Fig. 2B).

**Fig. 2.**
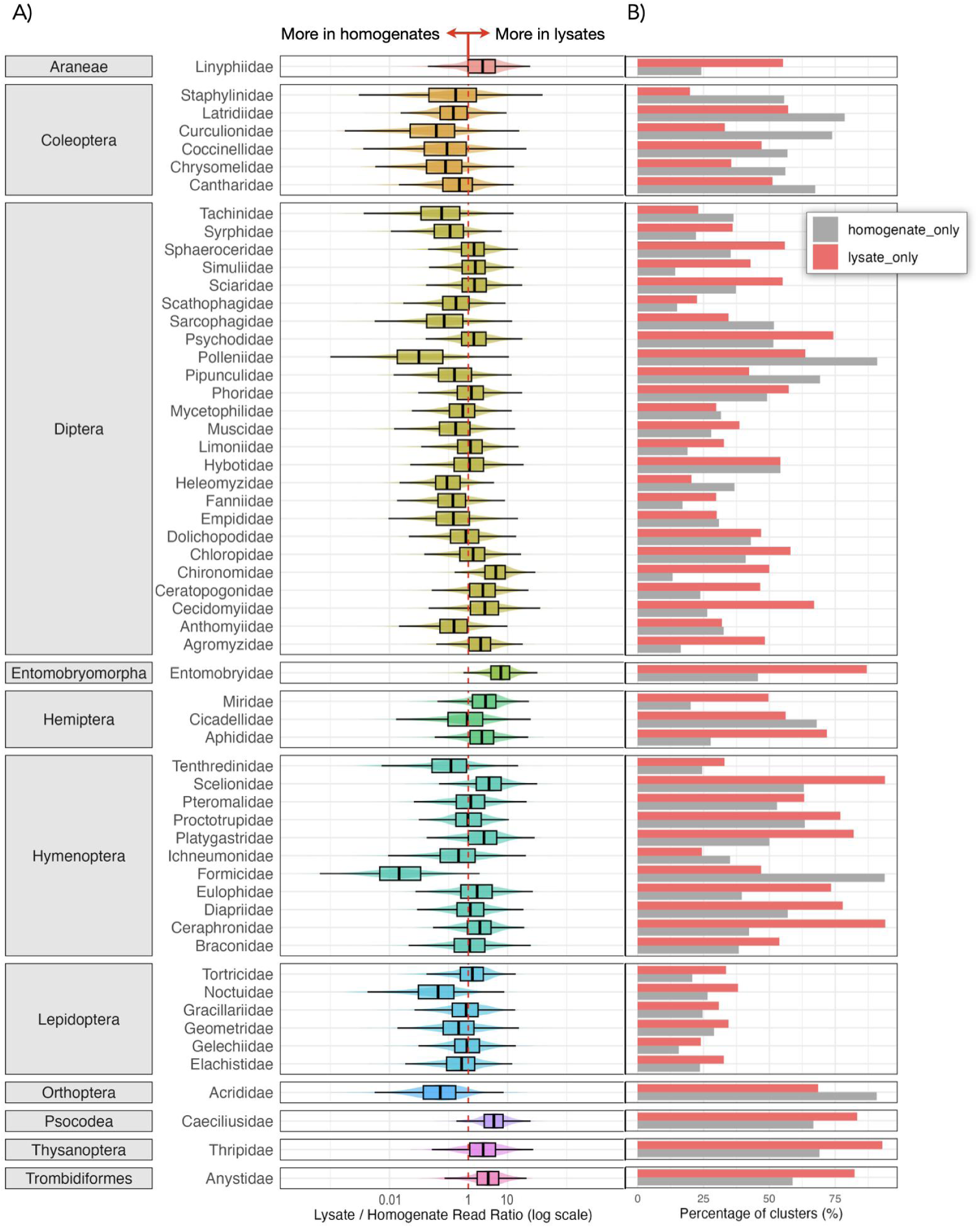
Influence of sample treatment on capturing arthropod diversity. **(A) Lysate-to-homogenate read ratios for 856 treatment pairs.** Data are aggregated by family, grouped by order. The violin plots show the distribution of treatment-pair ratios for each family and overlaid boxplots indicate the median and interquartile range. The red dashed vertical line marks a ratio of 1, indicating equal recovery from lysates and homogenates. Ratios < 1 (left) indicate higher recovery from homogenates, while ratios > 1 (right) indicate higher recovery from lysates. Only families occurring in at least 500 or more samples are shown. **(B) Percentage of arthropod clusters detected exclusively in either lysates or homogenates, aggregated by family.** The X-axis shows the proportion of clusters (out of all clusters detected in that family across both treatments) that were found only in one treatment. Bar colors indicate the treatment in which these clusters were recovered: grey – homogenates; pink – lysates.

### Individual insect barcoding

To compare metabarcoding data to data from individual barcoding, we selected 15 samples for which we performed both metabarcoding of lysates and homogenates *and* individual insect barcoding (Fig. 1C). These samples, hereafter referred to as *reference samples*, were collected at three sites in Southern Sweden (marked with diamonds in Fig. 1A) during five consecutive weeks in April and May 2019 (details in Table S1). Once these samples had undergone mild lysis, each individual was separately recovered and placed in a separate well on a microplate (Fig. 1C). When the specimen was too large to fit in the well, we used only one of its legs for barcoding, but also the remaining body during homogenization. To obtain DNA from these specimens, we adopted a HotSHOT procedure (Feng et al., 2026; Srivathsan et al., 2019; Truett et al., 2000). The resulting DNA was used as a template for individual insect barcoding, which involved amplifying a 313-bp fragment of the COI barcode for each individual using the m1COlintF and jgHCO2198 primers. Each primer was modified at 5’ end with 9-bp tags and a unique combination of tagged forward and reverse primers were used for specimen to sequence association (Geller et al., 2013; Leray et al., 2013; Meier et al., 2016; Yeo et al., 2021). Amplicon libraries were then prepared by ligating Illumina adapters to pooled products, and sequenced on Illumina MiSeq at 2×250bp flow cell at Genome Institute of Singapore.

Illumina PE reads were first assembled using PEAR v0.9.11 (Zhang et al., 2014) and data was demultiplexed using *ngsfilter* in OBITOOLs (Boyer et al., 2016) allowing for up to 2 mismatches in primer region and no mismatch in the tags. Quality and length of the pair-merged reads were assessed using *FASTQC v 0.11.9* (Andrews, 2010) and filtered using *VSEARCH v 2.22.1* (Rognes et al., 2016), with a minimum length set to 308 bp and maximum to 314 bp. Sequences were then dereplicated, singleton ASVs discarded, and the remaining sequences were ordered by descending number of reads. To account for erroneous sequences, including chimeras, we denoised each sample separately using the *UNOISE3* algorithm (Antich et al., 2021) implemented in the *USEARCH v 11.0.667* program (Edgar, 2010). The denoised amplicon sequence variants (ASVs) for all specimens were then clustered into operational taxonomic units (OTUs) based on 97% sequence similarity. Taxonomic labels were independently assigned to all ASVs using the same tool set as in HAPP (see above), and ASVs filtered to only retain those representing animal mitochondria. The chimeras were identified using the UNOISE3 algorithm integrated in USEARCH. Based on the resulting OTU-by-sample tables, we selected the most abundant ASV per individual to be its barcode. To accept this ASV, it had to be classified as an animal and have at least 20 reads, with those reads representing at least 60% of all non-chimeric reads from the sample.

### Matching metabarcoding results and barcodes

To link metabarcoding reads with individually sequenced barcodes, we performed *BLAST* searches (Altschul et al., 1990; Camacho et al., 2009, version 2.15.0) run locally in a Unix environment. The local *BLAST* database was constructed from all ASVs detected in the metabarcoding data for the 15 reference samples (8,345 ASVs), and the query dataset consisted of all unique barcode sequences obtained from individually processed specimens (1,035 haplotypes). A barcode sequence was considered a valid match only if it aligned to an ASV with 100% coverage and 100% identity (no mismatches, e-value close to 0).

### Spike-in calibration

The utility of spike-ins was evaluated in two ways. First, we used a simple linear model relating the number of reads of one of the spike-in species to the number of reads of the other spike-ins and the synthetic spike-ins (if present). This allowed us to use the full set of samples (856 for homogenates and 4,749 for lysates) to examine how effective the other biological spike-ins were (together, one at a time or in combinations of one to five) in controlling for sample-specific and species-specific variation in read numbers across samples for a chosen spike-in. For the homogenate samples, we could also compare the effect of using synthetic spike-ins, biological spike-ins or both to control for sample-specific and species-specific variation.

Specifically, let *r_tj_* be the number of reads of spike-in species *t* in sample *j*, and let *s_j_* be the total number of reads of synthetic spike-ins in sample *j*. Then the full model we explored has the form

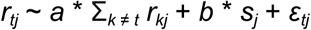

when using all remaining biological spike-ins for calibration, and the form

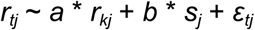

when using a particular spike-in *k*. Here, the factor *a* calibrates the read numbers against the sample-specific effect reflected by the biological spike-in reads, *b* does the same for the synthetic spike-in reads, and *ε_tj_* is the error term capturing the residual variation in *r_tj_*. The ratio of observed to fitted values can be used to measure the relative error that remains after calibration with spike-ins, and it can be compared with the ratio of the raw *r_tj_* values to their mean, reflecting the error in abundance estimates that would result from using the raw read numbers without calibration. By using different combinations of the *a* term and the *b* term, we assessed the effect of using biological spike-ins only, synthetic spike-ins only, or both types of spike-ins. In principle, each *r_tj_* value predicted based on spike-in data should be excluded from the fitting exercise for that prediction, but because of the large number of samples used, we assumed that the effect of excluding one data point could be safely ignored.

Second, we developed a hierarchical Bayesian approach to model the analysis steps more closely, potentially allowing more accurate estimates of species abundances. The model builds on our previous work, developed and tested using mock communities (Iwaszkiewicz-Eggebrecht et al., 2023). Here we take the same core Bayesian model, develop it further and test it on real insect communities (for a graphical representation, see Fig. S1). As the basis for developing and parameterizing the model, we used data from the 15 individually barcoded samples for which we held both metabarcoding results and information on the number of specimens included in the sample.

In brief, we model the number of reads *r* from sample *j* for species *t* with a Gamma distribution:

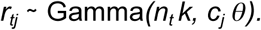

Here *c_j_*represents the “PCR factor“ per sample, *n_t_* is the number of insect specimens for species *t*, and *k* and 𝜃 define the shape of the distribution of available DNA strands per specimen that is available for PCR amplification.

The number of reads of the two synthetic spike-ins, *s_jt_*, was modeled in a similar way, with specific *k_s_* and 𝜃*_s_* values for the spike-ins, and *n* dropped:

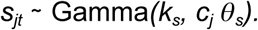

The model is based on the intuition that, if the distribution of DNA strands from a single specimen can be described by a Gamma(*k,* 𝜃) distribution, then the distribution from *n* specimens should be described by the sum of *n* independent variables from this distribution, which is known to be Gamma(*n k*, 𝜃). If we then multiply the number of copies with some (PCR) factor *c*, then this should be equivalent to scaling this gamma distribution, which is known to give us Gamma(*n k*, *c* 𝜃). Potentially, the *k* and 𝜃 values may differ between species, with the variation in *k* values reflecting the variation among species in the amount of DNA per specimen, and the variation in 𝜃 values reflecting the variation among species in PCR biases. There is an identifiability issue in this model between the PCR factor and the 𝜃 values, which we solve by fixing the average of 𝜃 values or the single 𝜃 value to 1.

As the objective of the modelling was to estimate abundances per species and sample from read counts, we adopted our earlier two-step approach (Iwaszkiewicz-Eggebrecht, Granqvist, et al., 2023): a training phase (step 1) and a prediction phase (step 2). In the training phase, the read numbers of spike-ins (biological, synthetic, or both) were used to train the model and to infer model parameters; in the prediction phase, the parameter distributions learned were used as input into a predictive step to infer the number of specimens (*n_tj_*) of the non-spike-in species from their read numbers *r_tj_* and the learned distributions of remaining model parameters. The predictive performance was assessed by comparing the estimates to the actual number of specimens in the sample.

The true value of *n* is known for the spike-ins, and is also constant per spike-in species across all samples. For all other species the number of specimens per sample*, n_j_,* is inferred in the prediction step with a geometric prior on *n*. The parameter *p* of this geometric distribution was drawn per species. Two different hyperpriors were explored for *p* (see Supplementary Fig. S2):

- An uninformative hyperprior: *p* is sampled from *a* Beta(1,9), allocating most of the mass to the interval of 1-20. This is similar to settings in our earlier work (Iwaszkiewicz-Eggebrecht et al., 2023), but weakly more informative by assigning more probability mass to *n*=1, which is common in the data, and by having no upper limit on *n*.
- An informative prior: By fitting species-specific p-values to the specimen counts in the reference data, a distribution of species’ p-values was compiled. The informative prior is a combined hyperprior that closely matches this reference data of *p*-values, using two parts: 30% probability of *p*=0.95, and 0.70% probability of drawing *p* from a Gaussian with mean of 0.4 and standard deviation of 0.25.

For the priors of the other parameters, we used a gamma prior on *k*, and a lognormal prior for both 𝜃 and *c*. The latter were assigned conjugate normal-inverse-gamma hyperpriors.

In step 1, the model was trained using three alternative training data sets: 1) only synthetic spike-in data (*s_j_* values; “synthetic spike-in models”); 2) only biological spike-in data (*r_tj_* values for the spike-in species; “biological spike-in models”); and 3) both synthetic and biological spike-in data (“combined models”). For the biological spike-in and combined models, we tested different combinations of shared or species-specific values of *k* and 𝜃 across spike-ins.

We implemented these different model versions in *TreePPL*, a probabilistic programming language aimed specifically at biological applications such as phylogenetics and biodiversity studies (Senderov et al., 2024). This language allows the user to define the model and then choose from a set of inference methods, allowing for easy comparison and exploration of both different model versions and inference strategies.

For step 1, both Sequential Monte Carlo (SMC) inference and Markov Chain Monte Carlo (MCMC) inference was used. SMC is an efficient way of calculating the normalization constants (logZ) per model, allowing model comparison using Bayes factors (Kass & Raftery, 1995). However, for some of the more complex model versions, the SMC algorithm did not fully converge, resulting in parameter estimates that were not satisfactory. Therefore, MCMC was used to estimate model parameters that were needed for step 2, predictions. In step 2, SMC was used with custom drift kernels per parameter, see the model scripts on github for separate kernel settings.

MCMC settings in step 1:

`-m ’mcmc-lightweight’ --kernel --mcmc-lw-gprob 0`

SMC settings in step 1:

`-m ’smc-apf’ --subsample --subsample-size 1 --resample manual`

SMC settings in step 2:

`-m ’smc-apf’ --subsample --subsample-size 1 --resample manual`

For convergence, we used the following convergence criteria: for SMC: the variance in the logZ over 50 runs should be less than 1.0; for MCMC, the parameter estimates should have an 𝑅^ below 1.01 across three independent runs starting from independent draws from the prior distribution. In step 1, all parameters of interest converged using the described MCMC approach. In the predictive step, all species converged apart from three clusters. These have been excluded from the figures.

## Results

### Effect of mild-lysis and homogenization on community recovery

Metabarcoding of the DNA obtained via mild lysis of 4,749 samples resulted in a median of 0.83 M filtered reads per sample (SD=0.44 M), forming a total of 33,989 clusters (proxy for species) in total. Metabarcoding of the subset also homogenized (856 samples) resulted in an average of 1.25 M reads per sample treated by mild lysis (SD=38,162) and 0.51 M reads for homogenates (SD=12,410). A total of 23,072 clusters were found. Of these clusters, 19,301 were shared between treatments, 1,302 clusters were found only in homogenates, and 2,469 were unique to lysates. Despite lysate samples being sequenced more deeply than homogenate samples (Fig. S3), we did not observe poorer cluster recovery from homogenate samples. The proportion of lysate clusters also recovered from homogenates did increase with sequencing depth (Fig. S4).

Arthropod families differed substantially in their relative representation in lysate vs homogenate samples (Fig. 2). Mild lysis resulted in better relative representation of many Diptera families, particularly small-bodied taxa such as Cecidomyiidae, Chironomidae, Ceratopogonidae, and Sciaridae. For these groups, many clusters were detected exclusively in lysates and not in homogenates (Fig. 2B; Fig. S5). In contrast, homogenization favored larger-bodied, more robust Diptera, including Polleniidae, Tachinidae, Syrphidae, Heleomyzidae, Fanniidae, Empididae, and Anthomyiidae. For these families, the difference was mainly quantitative, showing higher read counts in homogenates. It likely reflects the larger body sizes and stronger sclerotization of these taxa, while almost no clusters were unique to homogenates.

Among Hymenoptera, several families were more frequently detected in lysates. This included families Platygastridae, Ceraphronidae, Diapriidae, Scelionidae, and Eulophidae, all of which primarily consist of small-bodied parasitoids. Homogenates, however, yielded proportionally more reads from Formicidae (ants), although few clusters were unique to either treatment. Ichneumonidae, relatively larger and more heavily sclerotized, were also represented by more reads in homogenates, which additionally recovered unique clusters for this family. Overall, this suggests that homogenization enhances representation of ichneumonids in the metabarcoding data. In turn, Braconidae, one of the most abundant and diverse wasp families, showed a mixed pattern. Here, many clusters were shared between and represented in similar proportions in the treatments. However, some were found in one treatment alone, with lysate-only occurrences predominating (Fig. 2B).

Beyond Diptera and Hymenoptera, individual families in Entomobryomorpha, Araneae, Psocodea, Thysanoptera, and Trombidiformes were more often detected in lysates than in homogenates. Conversely, homogenization provided stronger representation for Acrididae (Orthoptera) and all Coleoptera families. These patterns corroborate the general inference that processing method influences species recovery in a lineage-specific manner.

### Individual specimen barcoding

After mild lysis, 4,804 individuals from the 15 reference samples were recovered. Amplicon sequencing for these individual insects resulted in barcodes for 4,173 individuals (∼87% success; details in Table S1; average 2,733 reads per individual; max = 27,128; min = 7). They belonged to 1,035 haplotypes of which 943 had a perfectly matching ASV in the metabarcoding dataset (91%; Fig. 3A, B). The matching haplotypes belonged to 474 metabarcoding clusters.

**Fig. 3.**
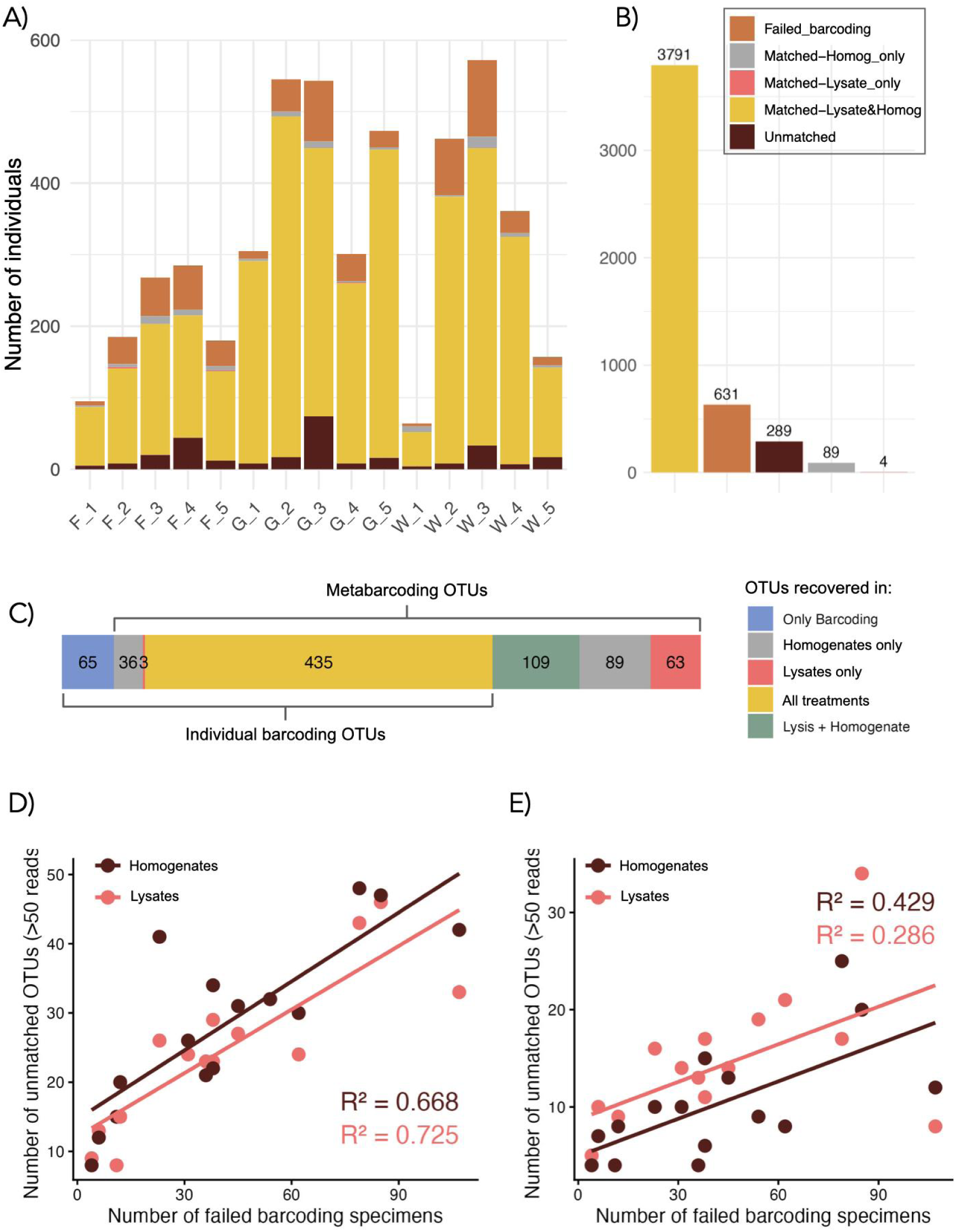
Summary of individual barcoding results and overlap with metabarcoding data. Panel **(A)** provides a histogram of specimen counts per sample. Colors indicate: failed barcoding (rusty orange), successful barcoding but not found in metabarcoding data (brown), and barcodes that successfully matched to metabarcoding data (remaining colors). This last category is subdivided into barcodes that are matched to both lysate and homogenate metabarcoding data (yellow), only lysates (pink) or only homogenates (grey). Panel **(B)** shows a cumulative summary of the same categories across all 15 samples. Legend applies to panels A and B. **(C)** Distribution of shared and split clusters (OTUs) between individual barcoding and metabarcoding treatments. **(D)** Number of individuals that failed barcoding per sample vs. number of OTUs recovered by metabarcoding but not matched to individual barcodes. **(E)** Comparison as in (D), with low abundance clusters (<50 reads per sample) filtered out.

For 289 specimens (92 haplotypes) representing 65 clusters, the ASVs of individually barcoded specimens had no matches in the metabarcoding data set (Fig. 3C). Of these, 72 specimens representing 26 clusters, were small parasitoids, flies/midges, microlepidoptera with only one leafhopper (genus *Hebata)* potentially representing a larger insect (Fig. S6). Those may have been missed during metabarcoding due to low biomass or primer mismatch. The remaining 217 specimens lacking a match in the metabarcoding data were mostly mites (N=212; 56 haplotypes belonging to 34 clusters). There were also one springtail (Collembola), two thrips (Thysanoptera), and two sequences that did not match any animal. Secondary signal with a strength of >20 reads could be recovered for 94 of the 212 specimens. It corroborated the main signal in all cases, suggesting that the specimens were correctly identified (Table S2).

Clusters found in metabarcoding but without a match in individual barcoding constituted almost 30% (198 out of 669) of the homogenate clusters and 28% (172 out of 610) of the lysate clusters (Fig. 3C). Despite those relatively high numbers of clusters, they were responsible for a minor proportion of total read counts (1.8% in lysates and 3.5% in homogenates; Fig. S7). The number of those metabarcoding clusters correlated well with the number of specimens that failed in individual specimen barcoding (Fig. 3D,E). This suggests that metabarcoding successfully detected many of the specimens that failed in individual-specimen barcoding. The low-read metabarcoding clusters were more common in homogenates as we see the number of clusters drop after filtering out low-abundance OTUs (Fig. 3D,E). Those clusters included obvious trace DNA from the environment (OTUs identified as cat, dog, squirrel, pheasant, perch, annelid worm etc), and DNA that could represent food items (springtails, Entomobryidae) (Fig. S8).

### Calibration using spike-ins

The raw read numbers of the six biological spike-in species varied widely across samples, despite identical numbers of individuals being added to each sample. This was true both for lysate (Fig. 4A) and homogenate samples (Fig. 4C), with the variance being considerably larger for lysates. Thus, abundance estimates based on raw read numbers without calibration would be associated with considerable error, particularly for lysates.

**Fig. 4.**
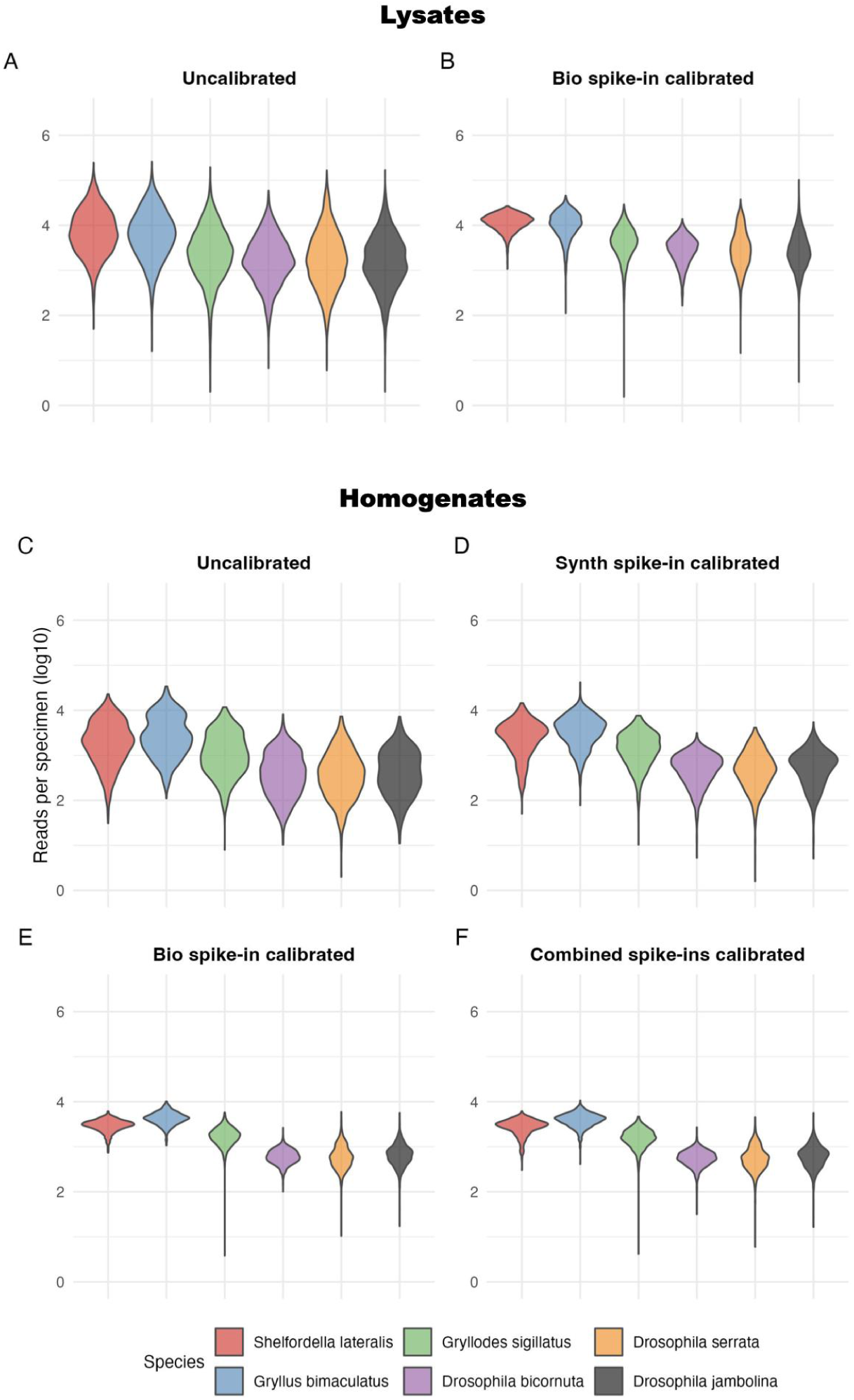
Calibration of read counts per specimen using biological or synthetic spike-ins to account for sample-specific effects. Log10-transformed read counts per specimen for six spike-in species (Shelfordella lateralis, Gryllus bimaculatus, Gryllodes sigillatus, Drosophila bicornuta, D. serrata, D. jambulina) across insect community samples. Panels (A and B) show lysate data (n=4490), and panels (C–F) show homogenate data (n=856). (A, C) Raw read counts. (B, D–F) Read counts after spike-in–based calibration. Panels (B, E) show calibration using biological spike-ins (each species calibrated against the other five). Panel (D) shows calibration using synthetic spike-ins (available only for homogenates). Panel (F) combines biological and synthetic spike-in calibration.

Applying spike-in calibration markedly reduced variation in the number of reads per specimen across samples. In lysates, the effect was particularly strong for *Shelfordella lateralis*, *Gryllus bimaculatus* and *Drosophila bicornuta* (Fig. 4B). A similar pattern was observed for homogenates, with even stronger variance reduction (Fig. 4E).

Calibration using synthetic spike-ins alone (only possible for homogenized samples), also reduced the variance in read counts per specimen across samples (Fig. 4D). Nonetheless, the improvement achieved was much slighter than the improvement achieved through calibration of homogenate data using only the biological spike-ins (Fig. 4E). Combining both spike-in types in calibrating homogenate counts (Fig. 4F) did not notably improve the results, as compared to using only biological spike-ins alone. These differences between calibration methods remained unchanged when the analysis was restricted to the 856 samples for which we had access to both lysate and homogenate data (Fig. S9).

The relative error (measured as the ratio of the actual read numbers per specimen to the fitted values after calibration with biological spike-ins) approximately followed a lognormal distribution (Fig. S10 and S11). Here, the standard deviation of the logarithm of the relative error (denote it *f*) gives an idea of the accuracy of the resulting abundance estimates; the interval (e/*f*,*ef*) would contain values within one standard deviation up or down from an estimate *e*. For lysates, using 5 spike-in species for calibration resulted in a standard deviation corresponding to a factor *f* of approximately 2.1 to 2.9, while for homogenates, *f* was in the range from 1.5 to 1.9 (Fig. S12). The number of spike-in species used for calibration had a positive effect on the accuracy. However, this effect leveled off after 3 to 4 species, indicating that it would be difficult to improve the accuracy much beyond what was achieved with the 6 spike-in species used in this dataset (Fig. S12).

Focusing on pairwise comparisons, the spike-in with the largest proportion of reads (*Sherfordella laterialis*) proved the most effective in reducing the variance in abundance estimates of the other species in lysates (Fig. 5B). In homogenates, the effect of the read proportion was less pronounced but the trend appeared to be the same (Fig. 5A). *Drosophila serrata* had the second largest proportion of reads, and reduced the variance in abundance estimates the most. Taxonomic affinity was a poor predictor of calibration efficiency: the best spike-in for a *Drosophila* species was often an unrelated species with large read count rather than another *Drosophila* species. The patterns observed in the full set of samples (4490 lysates and 856 homogenates) were consistent with those observed in the subset of samples for which we had both lysate and homogenate data (n=856).

**Fig. 5.**
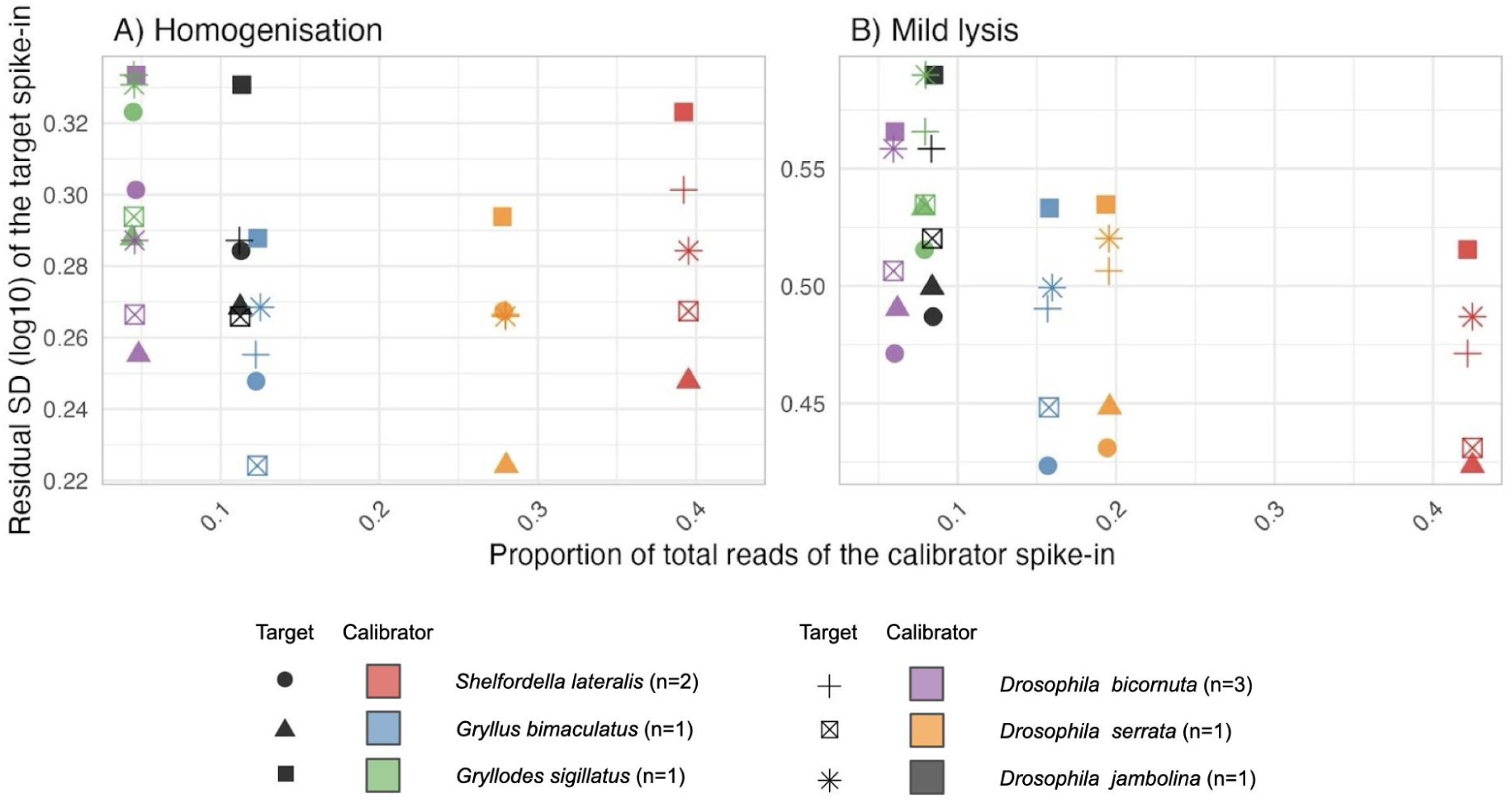
Pairwise calibration efficiency of all combinations of the six biological spike-ins. Each point represents a calibration of a target spike-in (marked in point shape) by using a calibrator spike-in (marked by color). We relate the proportion of total reads assigned to each calibrator spike-in (x-axis) with the residual SD of the target spike-in (y-axis) across samples treated by (A) homogenisation and (B) mild lysis. A lower residual SD indicates that the calibrator has worked better than for higher SD’s, as more of the variation across-samples was removed.

Finally, to assess the utility of calibration in practice, we explored the relationship between calibrated read counts and actual specimen numbers for non-spike-in species in the 15 reference samples. In these analyses, we focused on the species occurring in ten or more of the samples. For five of the six species satisfying these criteria, the calibrated homogenate read counts explained a large fraction of the variance in specimen numbers (R^2^ > 0.85; Fig. 6). The same trend was observed for only two out of six species for lysate read counts (Fig. S13), indicating that lysate read numbers do not reflect specimen numbers well for many species. These analyses also indicate that there are some species for which it will be challenging, if possible at all, to estimate abundance from read numbers even when focusing on homogenate data (Fig. 6B, *Polietes nigrolimbata*, a muscid fly).

**Fig. 6.**
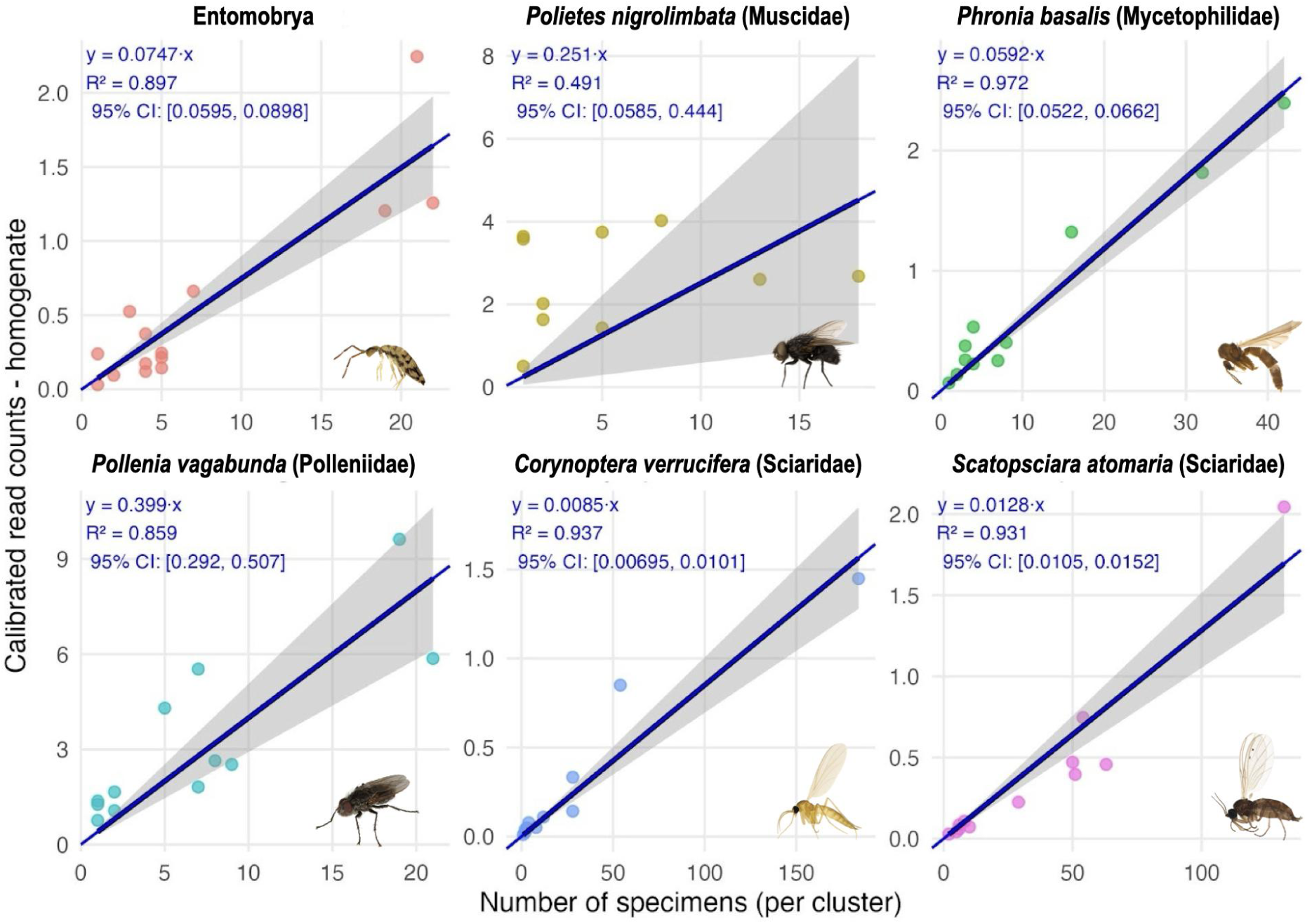
Relationship between calibrated read counts from metabarcoding (y-axis) and the number of specimens per cluster/species in the sample, as determined metabarcoding (y-axis). Included are data from homogenates of the reference 15 samples, with one panel for each of the six clusters occurring in 10 or more samples.

A potential concern with the biological spike-ins used here is that they are added to the samples in the laboratory, which means that they do not capture the variation among samples in the conditions experienced before reaching the lab, e.g., in the exposure to sun and extreme temperatures in the field. To assess the extent to which addition of spike-ins before field collection might improve the calibration, we compared the variance in calibrated reads per specimen of the spike-in species (solely reflecting variation arising during lab processing) versus the six non-spike-in OTUs occurring in ten or more of the reference samples (including also variation arising before samples arrived in the lab). This comparison suggests that the biological spike-ins do capture the major sources of variation in read numbers across samples (Figs. S14, S15).

### Abundance estimates from hierarchical Bayesian modelling

The Bayesian model comparisons showed that, regardless of which combination of spike-ins used, it was important to accommodate variation in PCR amplification factors (*c* values) across samples, at least when other sources of information were also accommodated properly (Fig. S16). Assuming a single PCR factor across all samples (1*c*) was never the best model.

For synthetic spike-ins, it was important to differentiate between the two spike-ins either in *k* values (amount of DNA) or in 𝜃 values (amplification bias), with a slight preference for the latter (Fig. S16A). Models accommodating variation in both did not improve model fit beyond this.

For biological spike-ins, it was similarly important to differentiate between species in either *k* or 𝜃 values, with a preference for models that differentiated among *k* values (amount of DNA per specimen; Fig. S16B). Again, models accommodating variation in both of these parameters did not improve model fit. When using both synthetic and biological spike-ins (Fig. S16C), we had difficulties estimating the normalization constant accurately with the SMC algorithm, but the results did indicate that the model with a shared 𝜃 value but eight spike-in-specific *k* values (six for the biological and two for the synthetic spike-ins) fit the data best, followed closely by the model with three *k* values (one for the biological and two for the synthetic spike-ins).

In the predictive step, we explored calibration models that used only the biological spike-ins and those that combined biological and synthetic spike-ins. In both cases, we compared the effect of ignoring or accommodating species-specific differences among biological spike-ins in DNA release (using species-specific *k* values). When using each of the six biological spike-in species as the hypothetical target for estimating specimen numbers, the calibration models that differentiated among the biological spike-ins clearly did best (Fig. S17). Including the synthetic spike-in data had only a minor effect on the results: they improved the estimates slightly for *Drosophila bicornuta* and *Gryllodes sigillatus*, but made them less accurate for *D. serrata* and *Gryllus bimaculatus*. As one might predict, the posterior distributions were weighted towards the biological spike-in distribution of the correct species, that is, the analysis was able to find the best matching calibration curve.

Finally, the number of specimens of the full set of non-spike-in species in the 15 reference samples was predicted using the “match 6*k*” model and both the uninformed and informed prior settings (See Fig. S2 for priors). The informed prior (Fig. 7A, orange bars) is shaped by the expected frequency of different specimen counts, that is, it is biased towards predictions of one specimen, as we know before seeing the data that the specimen count is likely to be small. The uninformed prior spreads out the prior probability more evenly on different specimen counts (Fig. 7B, pink bars).

**Fig. 7.**
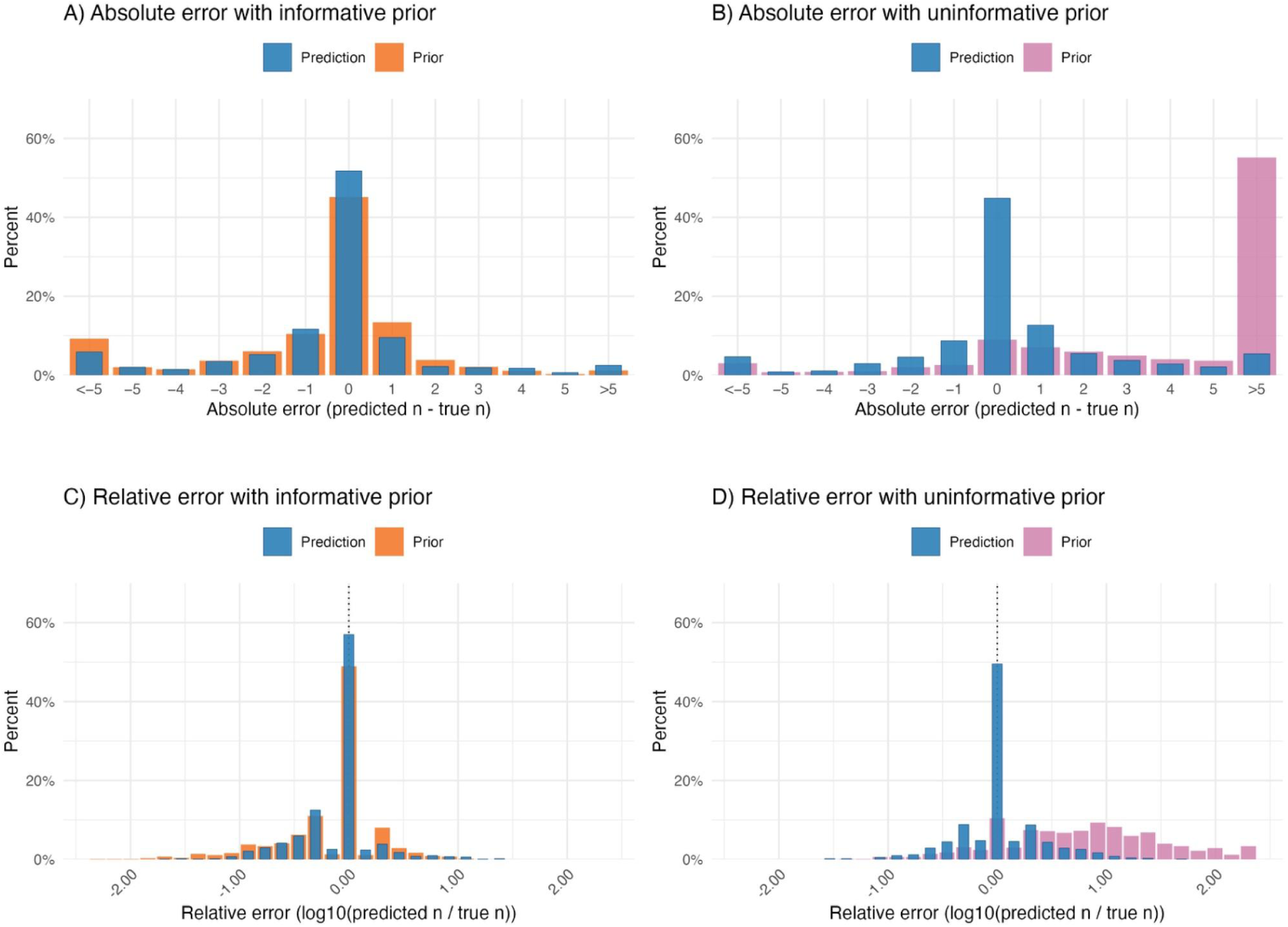
Prediction errors for the complete set of species’ occurrences for the fifteen samples, homogenates, with both the informed and uninformed hyperprior settings. The total number of (non-spike-in) species across the 15 samples was 465. Only three of these did not converge under the informative prior and were therefore excluded from the histogram. Under the uninformative prior, all 465 species converged. Error in the top panel is absolute error, calculated as the predicted values (the mode of the SMC sweeps, weighted by the normalization constant for each sweep) minus the known number of specimens (from individual barcoding). Error in the bottom panel is relative error, calculated as the log10 of the ratio between predicted values (the mode of the SMC sweeps, weighted by the normalization constant for each sweep) and the value from the reference sample. The “match model 6k” was used for all predictions.

Estimates of the number of specimens in the full set of non-spike-in species in the 15 reference samples—based on the biological spike-ins and accommodating species-specific differences in DNA release—were accurate (the absolute prediction error being 0 or +/- 1 specimen) in 72.9% of the cases under the informative prior, and in 66.3% of the cases under the uninformative prior. The metabarcoding data are clearly informative about the abundances under both priors, dramatically reducing overestimates under the uninformative prior and improving the precision under both priors.

## Discussion

This study addresses a key gap in the development of robust, quantitative biodiversity monitoring based on DNA metabarcoding. In summary, our results show that metabarcoding data can yield informative estimates of both species-specific occurrence and abundance in complex environmental samples of insect communities containing thousands of specimens and hundreds of species.

Our results document the precision and accuracy of occurrence data from the two dominant protocols, mild lysis (non-destructive) and homogenization (destructive). They also illustrate the uncertainty one can expect in abundance estimates from metabarcoding data using spike-ins. Specifically, we addressed four key methodological questions using a combination of large-scale Malaise trap data, individually barcoded samples, spike-in calibration, and Bayesian abundance modelling. In the following, we discuss our results in relation to each of these questions.

### How does mild lysis and homogenization affect community recovery?

Our results demonstrate that the majority of arthropod biodiversity in Malaise trap samples is captured by both treatments - mild lysis and homogenization - with 84% of taxa shared between treatments. However, important taxon-specific differences remain, reflecting both body size and structural traits of the taxa. Mild lysis recovered a greater number of small and soft-bodied taxa that were often no longer detected after homogenization. These included, among others, small hymenopterans such as Platygastridae and Ceraphronidae, as well as several Diptera families including Cecidomyiidae, Chironomidae and Sciaridae. Notably, all of these groups rank among the globally most abundant and diverse families (Santos et al., 2025; Srivathsan et al., 2023), and also among the least explored. Capturing this diversity is therefore particularly important in biodiversity surveys, as it represents a substantial and often overlooked component of global insect diversity. In addition, mild lysis enables the detection of these taxa while keeping specimens largely intact and available for further study. Once such taxa are detected through metabarcoding, researchers can return to the preserved samples for targeted sorting, morphological examination, or additional DNA-based analyses focused on the insects of interest (Feng et al., 2026; Iwaszkiewicz-Eggebrecht et al., 2025), or their biological interactions, including with microorganisms or parasites (Andriienko et al., 2024). This dual advantage—comprehensive detection combined with specimen preservation—underscores the value of non-destructive approaches for integrative biodiversity assessment.

Homogenates, on the other hand, yielded relatively higher total read counts than mild lysis when it came to robust-bodied taxa, including large Diptera (e.g. Pollenidae, Tachinidae, Syrphidae), and Orthoptera. This pattern is consistent with expectations, as these groups contribute substantially more biomass to homogenized samples, leading to higher representation of their DNA in the homogenate. Strongly sclerotized taxa, such as Coleoptera and Hymenoptera (particularly Formicidae and Ichneumonidae) were also much better represented in homogenates, likely because of the limited release of DNA through their rigid exoskeletons during mild lysis. Although fewer species were detected exclusively in homogenates than in lysates, unique detections occurred among several coleopteran families, among Acrididae (Orthoptera), and especially among Ichneumonidae - all of which are also among the world’s top ten most diverse insect taxa (Santos et al., 2025). These results highlight that homogenization remains valuable for recovering heavily sclerotized taxa that may be underrepresented in non-destructive treatments. Nonetheless, this advantage comes at the cost of destroying specimens and thereby losing the opportunity for subsequent study of the morphology or genetics of individual specimens.

### How accurate are occurrence estimates from metabarcoding?

Our analysis shows that the vast majority of insect specimens in a Malaise trap sample are detected by metabarcoding, and thus support the recent results of Furneux et al, (2025). Disregarding apparent errors in single-specimen barcoding, our results indicate that as much as 93% of the specimens in the samples were discovered by metabarcoding. The specimens that were missed in metabarcoding mostly represent small or low-biomass taxa, with a clear overrepresentation of mites (Acari) and parasitic wasps (Hymenoptera), suggesting that primer mismatch may be partly to blame. Hymenopterans are known to fail more frequently in single-specimen barcoding than other insects (Hebert et al., 2016), consistent with there being more variation in COI primer sites in this order than in other insects. Furneaux et al. (2025) report low detection rates for mites in DNA metabarcoding of Malaise trap samples using mild lysis and primers similar to ours. They speculate that this is due to the tough shell of mites releasing little DNA. However, we found that the mite detection rates were low in both lysates and homogenates, suggesting that primer mismatch is the more likely cause. In metabarcoding we used a different set of primers from individual barcoding and the mites did not seem to have amplified well with it.

Our results suggest that metabarcoding was successful in picking up signal from many of the specimens that failed in single-specimen barcoding (Figs. 3D, E). Metabarcoding also detected signal from a number of taxa that clearly represented trace DNA from the environment, and probably also from OTUs that represented food items. The latter OTUs were generally present in low read numbers, consistent with the results reported by Furneaux et al (2025). Homogenates appeared to pick up more of the trace DNA signal than lysates, suggesting that crushing the insects in the samples released more of the DNA from associated taxa, such as food items in the gut.

The 856 samples for which we had results from both lysates and homogenates revealed interesting OTU detection patterns beyond the general trends discussed previously (Supplementary Figs. S18, S19): For instance, lysates frequently failed to detect some of the largest, most sturdy insects, including bumble bees, some large flies and some beetles – even though these insects tend to be reliably detected in large read numbers in homogenates. Second, lysate read numbers vary more than homogenate read numbers, with low read numbers being more frequent. Specifically, the “tails” of the violin plots of Figs. S18–S19 often exhibit a slight widening around low read numbers in lysates, but rarely do so in data from homogenates. The fact that homogenates retrieved more “spurious” clusters that were not matched in individual-specimen barcoding than lysates (Fig. 3C) suggests that the wide tails largely reflect increased variability in DNA release among specimens in lysates.

Although homogenate samples were sequenced more shallowly than lysates (Fig. S3), sequencing depth had little effect on the detection of lysate clusters in homogenates (Fig. S4), indicating that the observed patterns are unlikely to result from insufficient sequencing. This also suggests that increasing sequencing depth alone will have limited success in recovering very small or low-biomass taxa from homogenates, and that alternative approaches such as size fractionation may be more effective, despite the additional logistical requirements (Ascenzi et al., 2025; Elbrecht et al., 2021).

In conclusion, metabarcoding of both lysates and homogenates tend to detect most insect species, and the species missed tend to be fairly strongly biased towards certain taxa, with Hymenoptera as the prime example. Such biases may be possible to address by further fine tuning of the primers, or the use of primer cocktails (Elbrecht et al., 2019). Lysates detect more OTUs on average, but this surplus may at least in part be due to increased chances of detecting trace DNA from past meals and interaction partners (see Furneaux et al. 2025).

### Can abundance estimates be improved with spike-ins?

Accurately estimating species abundance (in terms of biomass or specimen numbers) from metabarcoding data remains one of the major challenges in molecular biodiversity assessment (Elbrecht & Leese, 2015; Harrison et al., 2021; Luo et al., 2023). Clearly, sequencing read counts do not directly translate into species abundances, as they are affected by numerous biological and technical factors—including differences in body size, DNA content, extraction efficiency, PCR amplification bias, stochastic variation during library preparation, and sequencing depth (Krehenwinkel et al., 2017; Martoni et al., 2022). On the other hand, read counts should be informative about abundances given appropriate calibration and statistical analysis. In recent years, spike-ins have emerged as a promising means to calibrate read counts across samples and minimize technical noise (Harrison et al., 2021; Ji et al., 2020; Tourlousse et al., 2017).

The two types of spike-ins used so far – biological (standardized sets of insect individuals added to community samples) and synthetic (plasmids carrying artificial DNA constructs) – come with different pros and cons. Synthetic spike-ins can be designed appropriately so that they are not confused with real biological signal, they can be ordered cheaply in large quantities, and a precise amount of DNA copies can be added to each sample. Thus, they are quite convenient to use, and should provide effective controls for batch effects and sequencing biases. However, they are typically added relatively late in the process, and cannot control for variation among samples in the steps prior to DNA extraction.

By comparison, biological spike-ins (specimens of foreign species added to samples in predetermined numbers) may be more difficult and expensive to obtain. One may have to control for the variation among individual specimens in size or developmental stage, for instance by selecting specimens of a particular life stage and of “normal” size. The chosen species may later be found to occur naturally in the samples, or its barcode sequence may be confused with that of naturally occurring species. There is also a risk that the genome of the species generates off-target sequences, such as NUMTs, which can be confused for authentic metabarcoding signal. Another potential risk is that the chosen species is reared together with other species, whose DNA may accidentally be introduced as contamination with the biological spike-ins, and these species may also occur naturally in the samples.

Regardless of these challenges, there are also benefits that come with biological spike-ins. Most importantly, they can be introduced early on in the process, and capture more of the sources of variation affecting the read numbers per specimen. Theoretically, they can be added to sample bottles just before they are mounted on the traps in the field. However, this requires continuous access to suitable specimens close to the collecting sites. More realistically, biological spike-ins are added when the samples arrive in the lab. Even in the latter case, they capture variation among samples in lysis or homogenization conditions and in the efficiency of DNA extraction, in addition to all the subsequent steps that are covered by synthetic spike-ins.

Our results for homogenate data clearly indicate that the most important sources of variation in specimen read numbers stem from differences in the sample processing before DNA amplification and sequencing, that is, variation that is captured only by the biological spike-ins. As a result, biological spike-ins proved more effective than synthetic spike-ins in reducing variation in reads per specimen, and using both types of spike-ins did not perform any better than using biological spike-ins alone (Fig. 4). While we did not test synthetic spike-ins for lysate samples, there is no reason to believe that the result would have been different for these. In other words, it appears that biological spike-ins capture virtually all of the treatment variation among samples measured by synthetic spike-ins, and the potential increase in precision that synthetic spike-ins might bring does not appear to make a difference in practice.

When it comes to the choice of biological spike-in species, our results indicate that species that consistently yield larger read numbers are generally preferable (Fig. 5). The accuracy increases with the number of spike-in species but the improvement is small beyond three or four species (Fig. S12). It also appears that there is variation among spike-in species in how closely they match particular target species (Fig. 5). Interestingly, the most closely related species does not necessarily provide the best calibration of a particular target species. Thus, having access to a range of spike-in species with different responses to variation in treatment conditions appears beneficial.

Our results also demonstrate clear differences between lysate and homogenate treatments in the success of calibration. Variation in read numbers per specimen is considerably larger in lysates than in homogenates to start with, and the difference is even more pronounced after calibration (Fig. 4). The same pattern occurs in the 15 individually barcoded samples (Fig. 6). There are species for which it appears possible to obtain good abundance estimates from lysate read counts (one or two out of the six analyzed) but for the remaining species it appears to be challenging. The opposite can be said for homogenates; while read numbers correlate well with specimen numbers for most species (five out of the six), there seems to be species where this is not the case.

### How accurate abundance estimates can be obtained from metabarcoding data?

To obtain accurate estimates of the number of specimens, it would clearly be advantageous to develop a stochastic model that captures our understanding of the entire metabarcoding process. The hierarchical model we developed in this paper explicitly accounts for variation in amplification efficiency and other sources of technical and biological bias affecting read numbers of species occurring in the samples. In a Bayesian framework, we can use biologically informed priors and condition the inference of specimen counts of OTUs on the known data for biological and synthetic spike-ins. Thus, this analysis helps us translate raw read counts into posterior distributions on the likely specimen counts for the OTUs in the sample. It captures uncertainty while improving the interpretability of metabarcoding data, providing abundance estimates that more closely reflect the true underlying specimen numbers across diverse taxa.

The result is encouraging: validation against the individually barcoded reference samples, for which the true number of individuals per species was known, showed that the Bayesian model substantially improved the correlation between predicted and observed abundances compared to uncorrected read counts or prior expectations. Under an informative prior, the analysis provided spot-on estimates for 51.7% of species occurrences (a species occurrence refers to the specimen count of a particular OTU in a particular sample) and reasonably accurate ones (±1 individual) for 72.9% of occurrences.

Three observations from this analysis should be highlighted. First, the statistical estimation of species abundances in Malaise trap samples is a little bit similar to the problem of predicting the weather tomorrow. With good prior knowledge, it is easy to guess the right answer without having access to any data: the weather tomorrow is going to be the same as the weather today, and there is likely just one specimen of the species in the bottle. It is difficult for an analysis of empirical data to improve on such a prior; nevertheless, our abundance estimates under the informative prior did show a clear, if small, improvement. Potentially, the fact that the effect was so small could partly be due to the fact that we derived our informative prior from the actual samples used in the analysis. Nevertheless, our experience suggests that the prior we arrived at is fairly typical of Malaise trap catches in general. It is reassuring that our results under an uninformative (some might say “stupid”) prior show that there is enough information in the read counts of OTUs and spike-ins to generate reasonably accurate abundance estimates even under such adverse conditions.

Second, the species abundance distribution in Malaise trap catches seems to be a mixture of two underlying distributions. The majority of species are well characterized by a geometric distribution that is strongly skewed toward small numbers of specimens. However, some species seem to either be absent from the samples or occur in large numbers. This led us to use a mixture of geometric distributions as our informed prior. We speculate that the second group of species consist of those that tend to swarm in large numbers, or where the life stages that end up in Malaise traps are characterized by sharp spatiotemporal abundance peaks.

Finally, our analysis suggests that different species respond differently to the inevitable variations in the metabarcoding treatment across samples. For instance, models that separate the calibration curves for different spike-ins are strongly favored over models that assume they behave the same (Fig. S16). This is consistent with the observation from the simple calibration analyses, where the calibration curves of different spike-in species were compared among themselves (Fig. 5). We also noticed improvements in abundance estimates when allowing each OTU to “select” the best calibration curve among the biological spike-ins. These observations, taken together with the variation among species in the expected specimen counts alluded to above, strongly suggests that abundance estimates can be improved further by collecting calibration data for more species. Ideally, such data should be collected for all species for which one would like to obtain accurate abundance estimates. Of course, this will be difficult to achieve for rare species but should be possible for all species that are frequently encountered in monitoring programs.

### Towards an optimal workflow for Malaise trap and DNA-based biodiversity assessment

Our study reveals clear trade-offs among alternative metabarcoding approaches. Destructive and non-destructive methods differ in their sensitivity to particular taxa, as well as in costs, ease of application, and sample-storage requirements. Importantly, non-destructive methods preserve specimens for future study, whereas destructive homogenisation does not—but the latter yields data that permit more reliable abundance estimation. Solutions therefore need to balance cost-and work-efficiency while addressing multiple sources of variation.

For large-scale insect surveys and monitoring, an optimal strategy appears to combine both methods. The majority of samples should be processed using non-destructive lysis, which provides robust occurrence data and preserves material for targeted re-examination—an immense resource for future taxonomic and genomic research, if the storage demands can be met. A smaller subset of samples should be homogenized, given the improved ability to estimate abundances and recover taxa that may be missed by non-destructive approaches (e.g., endoparasites or weakly sclerotized specimens). For example, in weekly sampling schemes, homogenizing every fourth sample would enable tracking abundance trends while retaining most material for long-term investigation.

Another interesting way of eating your cake and having it too would be to adopt a two-step analysis workflow. First the samples are processed with mild lysis, so that undescribed taxa can be identified and recovered from relevant samples for description and single-specimen barcoding. When the new taxa have been properly documented, samples are homogenized to obtain good abundance estimates. As the reference libraries get increasingly complete, more and more samples will be completely characterized in the mild lysis step and can then go directly to homogenization.

Regardless of the approach chosen, and in addition to standard precautions (e.g., maintaining sufficient ethanol concentration during storage), our study underscores the importance of using spike-ins if abundance estimates are desired. Specifically, we can summarize our findings in terms of the following recommendations. If it is desirable to obtain abundance estimates, then homogenization should be preferred over mild lysis, and at least three or four biological spike-ins should be added to the samples as early as possible in the workflow. Ideally, this addition should happen just before the sample bottles are deployed in the field. Synthetic spike-ins do not markedly improve the calibration and can be skipped if biological spike-ins are used. When choosing biological spike-ins, species that consistently amplify well should be preferred, as well as sets of species representing a range of different body types and morphologies. Ideally, one should collect individual calibration data for all species for which accurate abundance estimates are desired.

Although commonly adopted in microbiome studies (Tourlousse & Sekiguchi, 2025), spike-ins have rarely been applied to arthropod diversity assessments so far (Iwaszkiewicz-Eggebrecht et al., 2024), despite their demonstrated value in translating relative read counts into more meaningful abundance estimates—a critical aspect of biodiversity monitoring. With AI-assisted taxonomic classification methods yet to achieve maturity, spike-ins currently offer the most practical means of bridging the gap between sequence data and specimen counts. They therefore represent an essential component of what may be considered the “holy grail” of metabarcoding: accurately reconstructing entire multi-species communities.

## Data and Code Availability Statement

Metabarcoding raw sequencing data is available at the European Nucleotide Archive (ENA) under study accession number PRJEB61109. Bioinformatically processed metabarcoding data is available at Figshare (https://doi.org/10.17044/scilifelab.25480681.v5) . Barcoding data will be made available on the NCBI’s SRA archive at the time of publication.

All scripts, including the *R* scripts and *TreePPL* model descriptions, as well as the data and *TreePPL* inference settings are available on github: https://github.com/insect-biome-atlas/paper-metabar-accuracy.git.

## Author contribution statement

**E. I-E.** conceived the project’s original idea; **E. I-E.**, **F.R.**, **P.L.**, and **R.M.** designed and developed the methodology. Metabarcoding was done with assistance from **M.B.** DNA barcoding was supervised by **R.M.** and **A.S.**, and the resulting data were analyzed by **C.V.** and **K.H.N. F.R.** and **E.G.** developed and implemented the model. Data analysis was performed by **E. I-E., F.R.** and **E.G.** Manuscript writing was led by **E. I-E.** and **F.R.** All authors contributed critically to the drafts and gave final approval for publication.

## Acknowledgements

This project was supported by the Knut and Alice Wallenberg Foundation (grant KAW 2017.088 to FR), the Swedish Research Council (grant 2021-04830 to FR), and the Polish National Science Centre (grants 2018/31/B/NZ8/01158 and 2021/43/B/NZ8/03376 to P.Ł.). The contribution of EG was supported by Mistra, the Swedish Foundation for Strategic Environmental Research, through the Finance to Revive Biodiversity program (grant DIA 2020/10). The contribution of TR was funded by the European Research Council (ERC) under the European Union’s Horizon 2020 research and innovation programme (grant agreement No 856506; ERC-synergy project LIFEPLAN). The data handling was enabled by resources provided by the Swedish National Infrastructure for Computing (SNIC), partially funded by the Swedish Research Council through grant agreement no. 2018-05973. Spike-in insect culture was supported by the National Institute of General Medical Sciences of the NIH under award number R35GM124701 to Brandon S. Cooper.

## Supplementary Materials for the manuscript Accuracy of occurrence and abundance estimates from insect metabarcoding

**Table S1.** Summary of the information concerning the collection of samples - habitat they were sampled from, ID of the trap, collection date, exact geographic location - and the values measured during pre-processing - preservative ethanol concentration (indicator of the quality of samples) and the wet weight of the sample (proxy for biomass).

| sample_ID |  | Habitat | trap_ID | collection_date | trap_collection_longitude | trap_collection_latitude | Ethanol conc. [%] | Wet weight [mg] |
| --- | --- | --- | --- | --- | --- | --- | --- | --- |
| STHFJP | G_1 | Grassland | TDGFXV | 13/04/2019 | 15.19485 | 56.93642 | 94.4 | 5.18 |
| S5MEZA | G_2 | Grassland | TDGFXV | 20/04/2019 | 15.19485 | 56.93642 | 94.4 | 5.00 |
| SY34ZS | G_3 | Grassland | TDGFXV | 27/04/2019 | 15.19485 | 56.93642 | 94.3 | 4.72 |
| SCNXZK | G_4 | Grassland | TDGFXV | 04/05/2019 | 15.19485 | 56.93642 | 94.8 | 1.27 |
| SKYQAL | G_5 | Grassland | TDGFXV | 11/05/2019 | 15.19485 | 56.93642 | 94.9 | 1.79 |
| SC1YIM | W_1 | Wetland | T7WIMF | 13/04/2019 | 14.75969 | 57.3824 | 95.1 | 1.42 |
| S8APX1 | W_2 | Wetland | T7WIMF | 20/04/2019 | 14.75969 | 57.3824 | 94.2 | 5.33 |
| SLWUQ2 | W_3 | Wetland | T7WIMF | 27/04/2019 | 14.75969 | 57.3824 | 94.4 | 4.47 |
| SIRUBH | W_4 | Wetland | T7WIMF | 04/05/2019 | 14.75969 | 57.3824 | 94.7 | 1.94 |
| SACBBR | W_5 | Wetland | T7WIMF | 11/05/2019 | 14.75969 | 57.3824 | 94.8 | 0.20 |
| SQVR9H | F_1 | Forest | TGXFSR | 13/04/2019 | 15.585445 | 58.28649 | 95.2 | 1.08 |
| SBWPUY | F_2 | Forest | TGXFSR | 20/04/2019 | 15.585445 | 58.28649 | 95.1 | 0.88 |
| SUXEKB | F_3 | Forest | TGXFSR | 27/04/2019 | 15.585445 | 58.28649 | 95 | 1.27 |
| SG3CB2 | F_4 | Forest | TGXFSR | 04/05/2019 | 15.585445 | 58.28649 | 95.1 | 1.45 |
| SIUQ3G | F_5 | Forest | TGXFSR | 11/05/2019 | 15.585445 | 58.28649 | 95.1 | 1.28 |

**Fig. S1:**
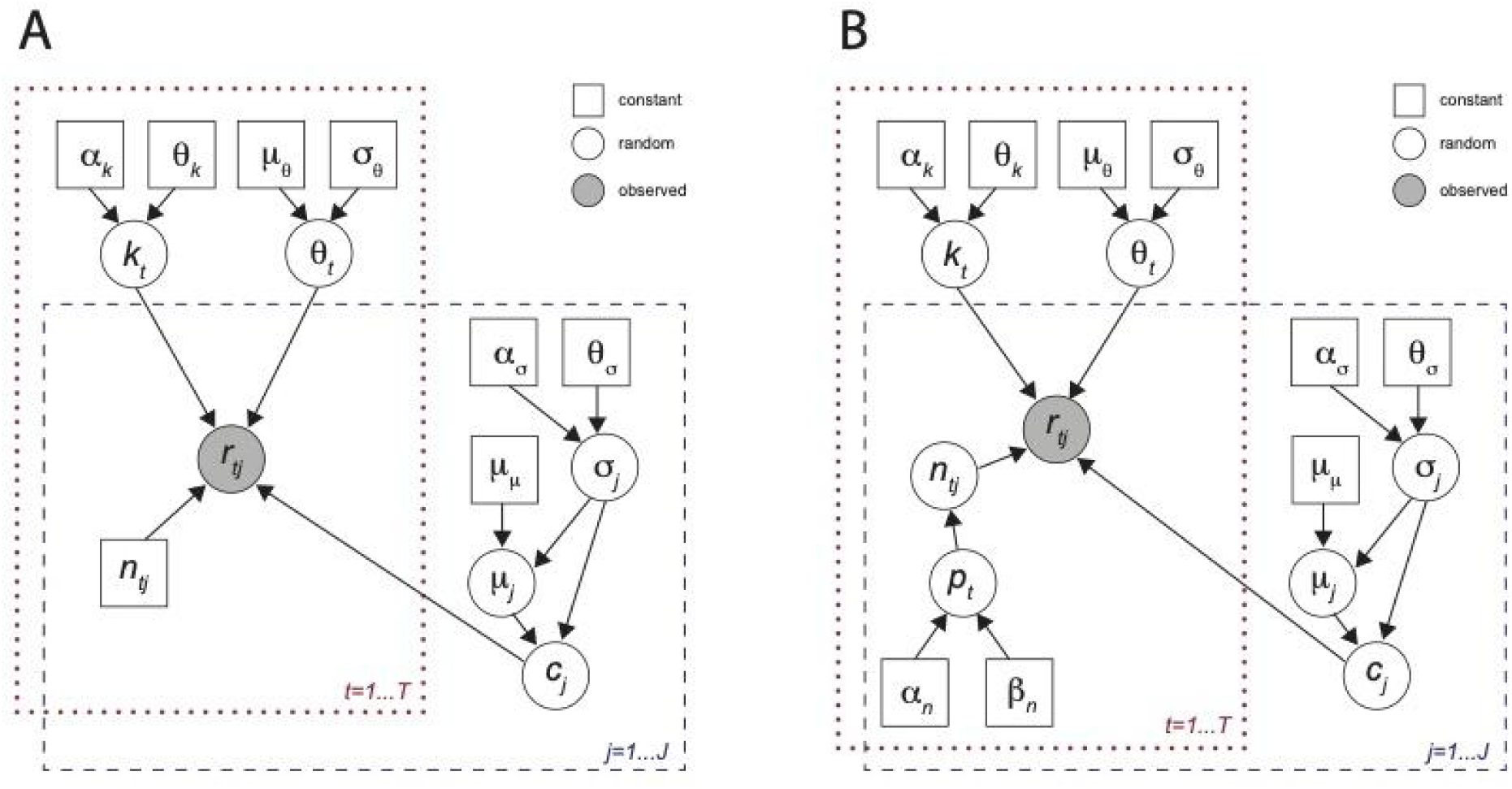
Probabilistic graphical model of abundance estimates model. r: reads counts, t: index for species, j: index for samples. Figure A shows step 1, where the posterior distributions of k, theta and c are learned and n are known constants (the specimen number per spike-in species. In B, step 2 is shown: the learned distributions from step 1 are used and n is inferred. The uninformative hyperprior for p is shown.

**Fig. S2:**
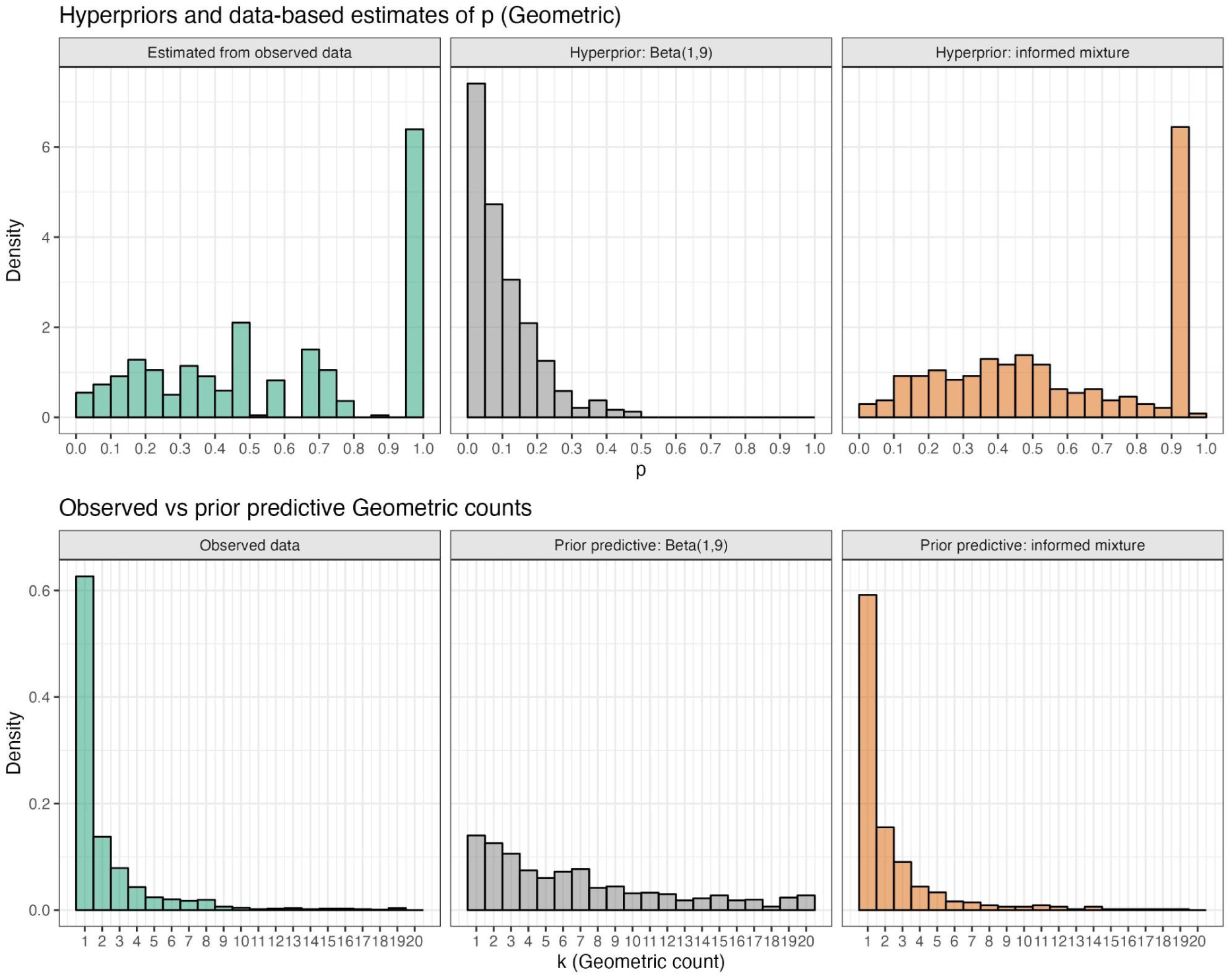
The top panel shows, from left to right, the estimated p-values per species from the reference data of the 15 samples (green), the “uninformative” hyperprior Beta(1,9) for p (grey) and the “informative” hyperprior on p which is a the mixture prior created to fit the estimated p-values per species from the reference data. The bottom panel shows the frequency of specimens per sample in the observed data (green), simulated using the uninformative hyperprior (grey) and simulated using the informed prior (orange). In the bottom panel, the x-axis has been limited to 1-20 to focus on the area of highest probability. In the observed data, the highest observed frequency value is 184.

**Figure S3.**
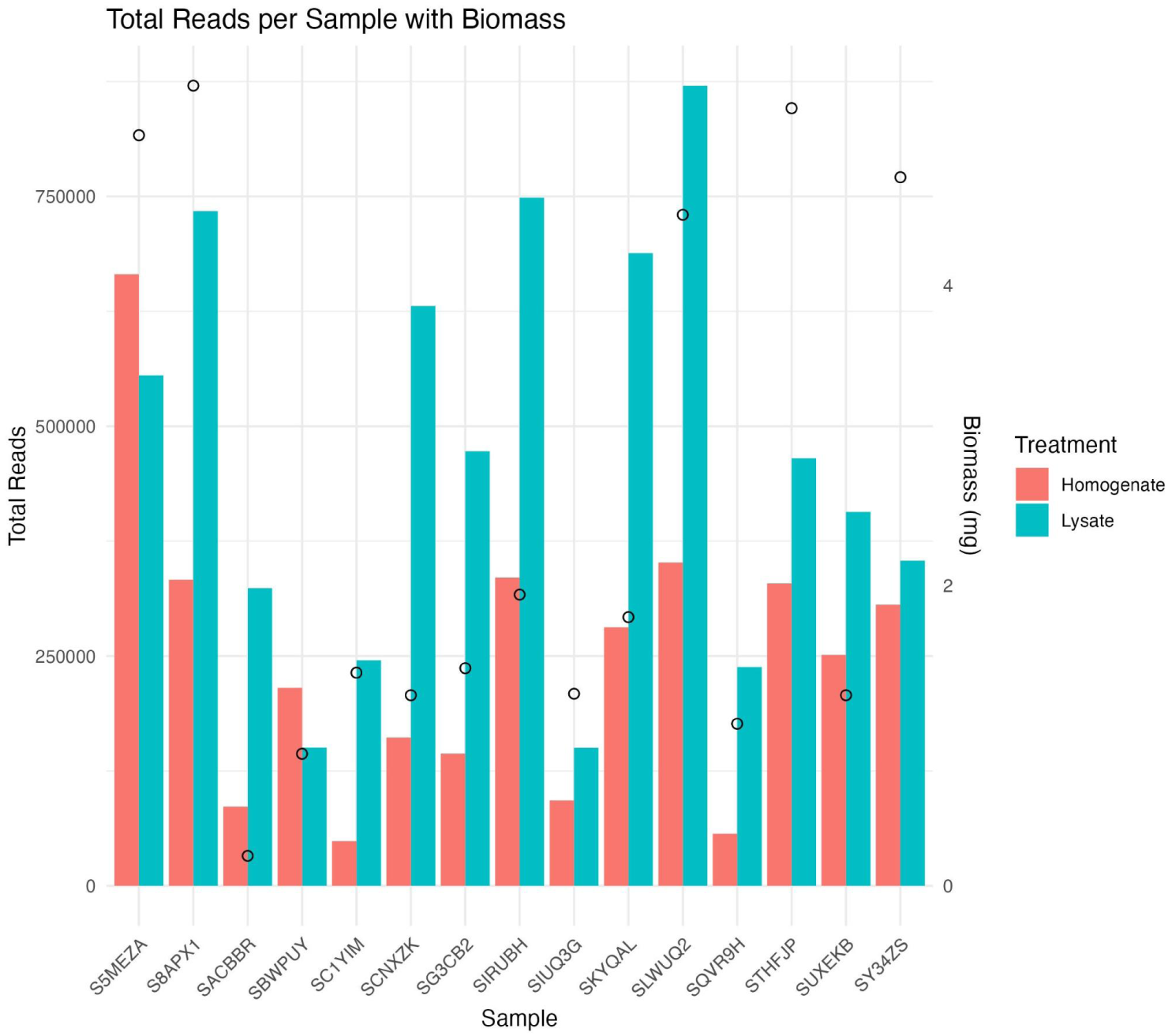
Selected samples - sequencing depth and biomass. Histogram shows the total number of reads per sample for both lysate and homogenate as well as the biomass (wet weight) of each sample. On average, homogenates received ∼50% of the reads of the lysates.

**Fig. S4.**
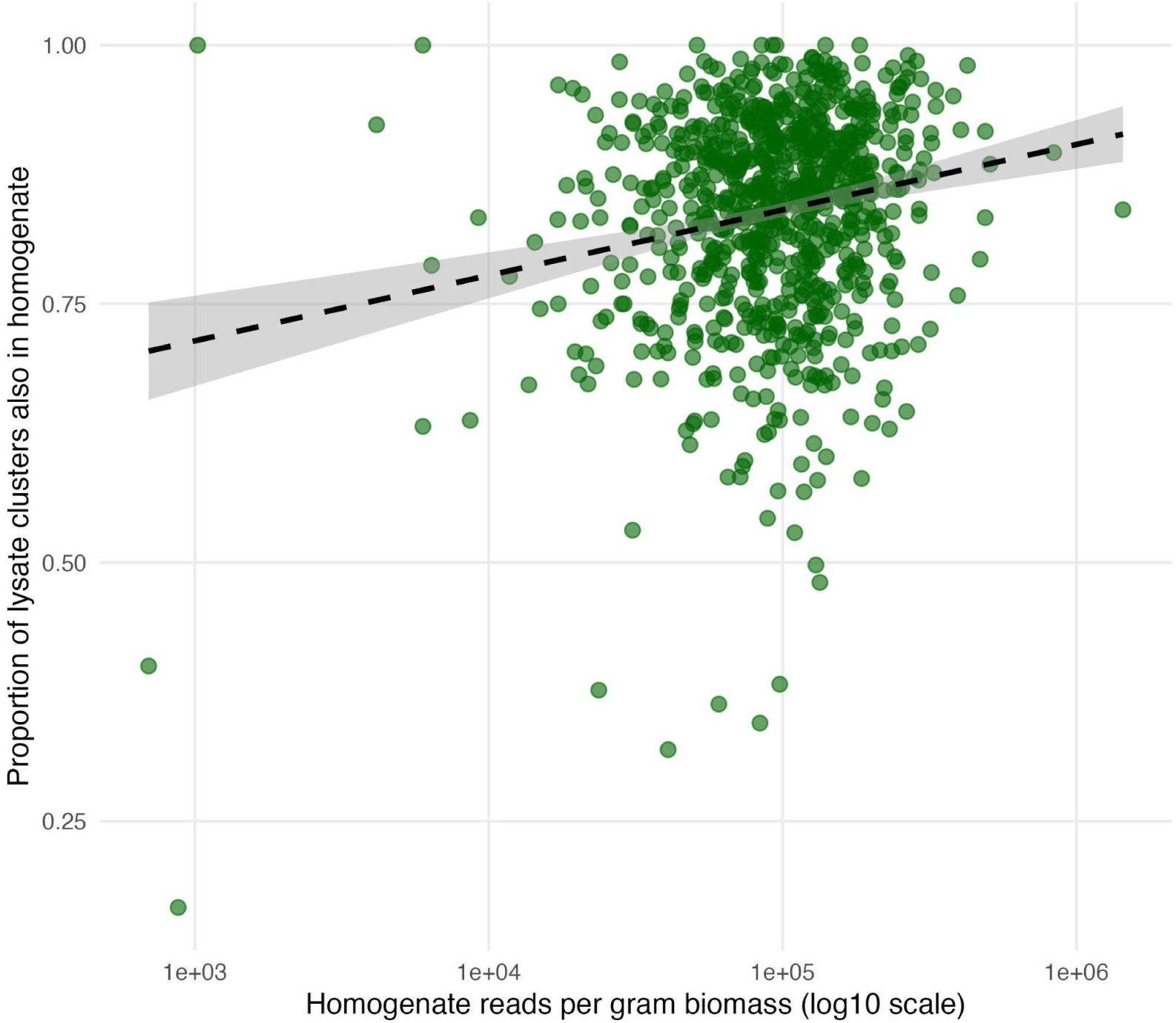
Relationship between the proportion of lysate clusters also retrieved in homogenates (y-axis) and the total number of homogenate reads per gram of biomass (x-axis). Each point represents one of the 856 Malaise trap samples that underwent mild lysis and homogenization. A dashed line indicates the linear regression fit with 95% confidence interval. Three samples with extremely low sequencing coverage (total homogenate reads per biomass < 100) were excluded to improve visualization.

**Fig. S5.**
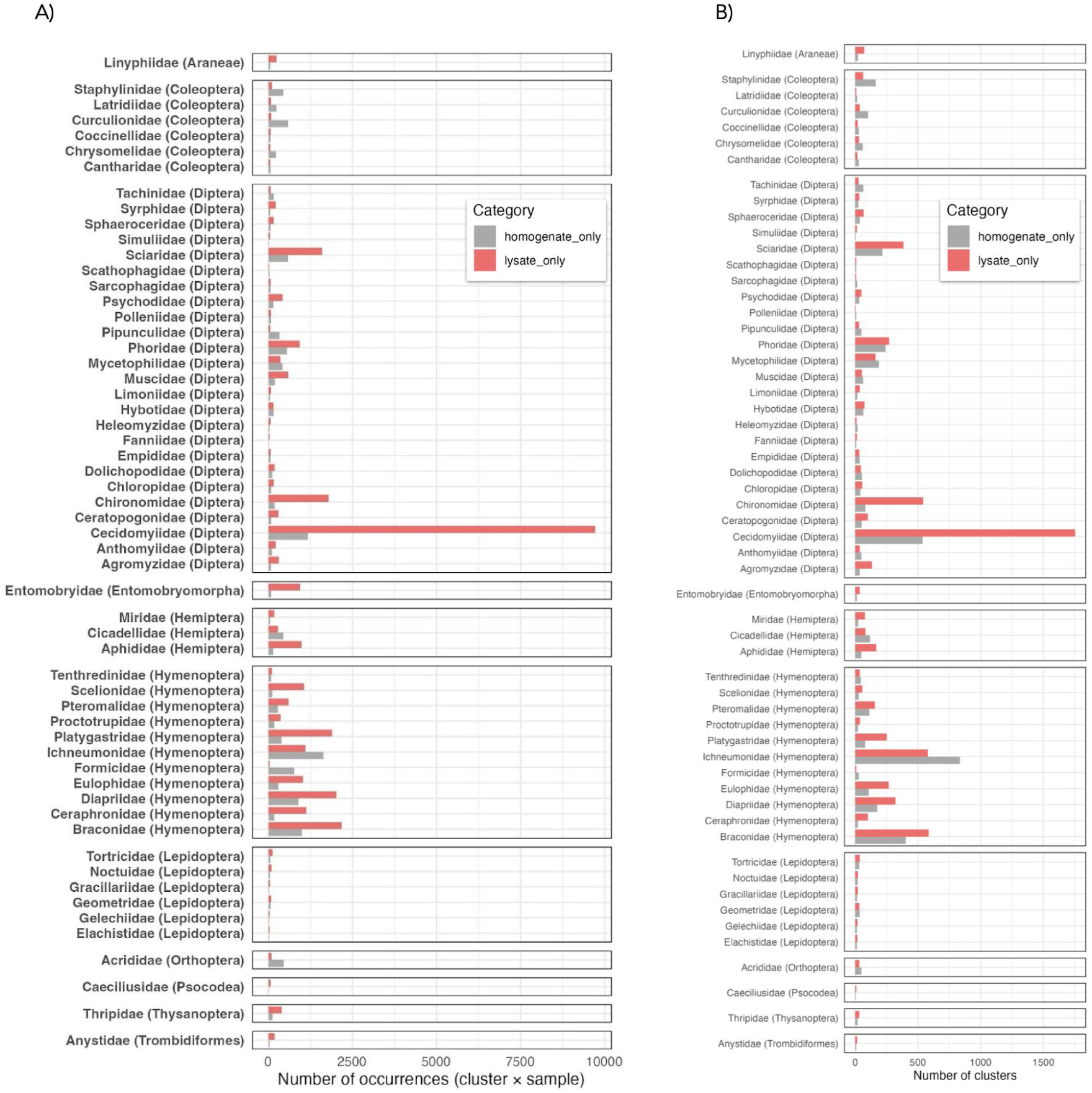
Influence of sample treatment on capturing arthropod diversity - number of instances of clusters being detected exclusively in either lysates or homogenates, aggregated by family. Arthropod clusters detected exclusively in either lysates or homogenates, aggregated by family. **(A)** The X-axis shows the number of times (samples) that a cluster was detected in one treatment but not the other. The color of the bar reflects in which treatment were those clusters recovered: grey - homogenates; pink - lysates. **(B)** The X-axis shows the number of clusters that were detected in one treatment but not the other (irrespective of the number of samples it occurred in). The color of the bar reflects in which treatment those clusters were recovered: grey - homogenates; pink - lysates.

**Fig. S6.**
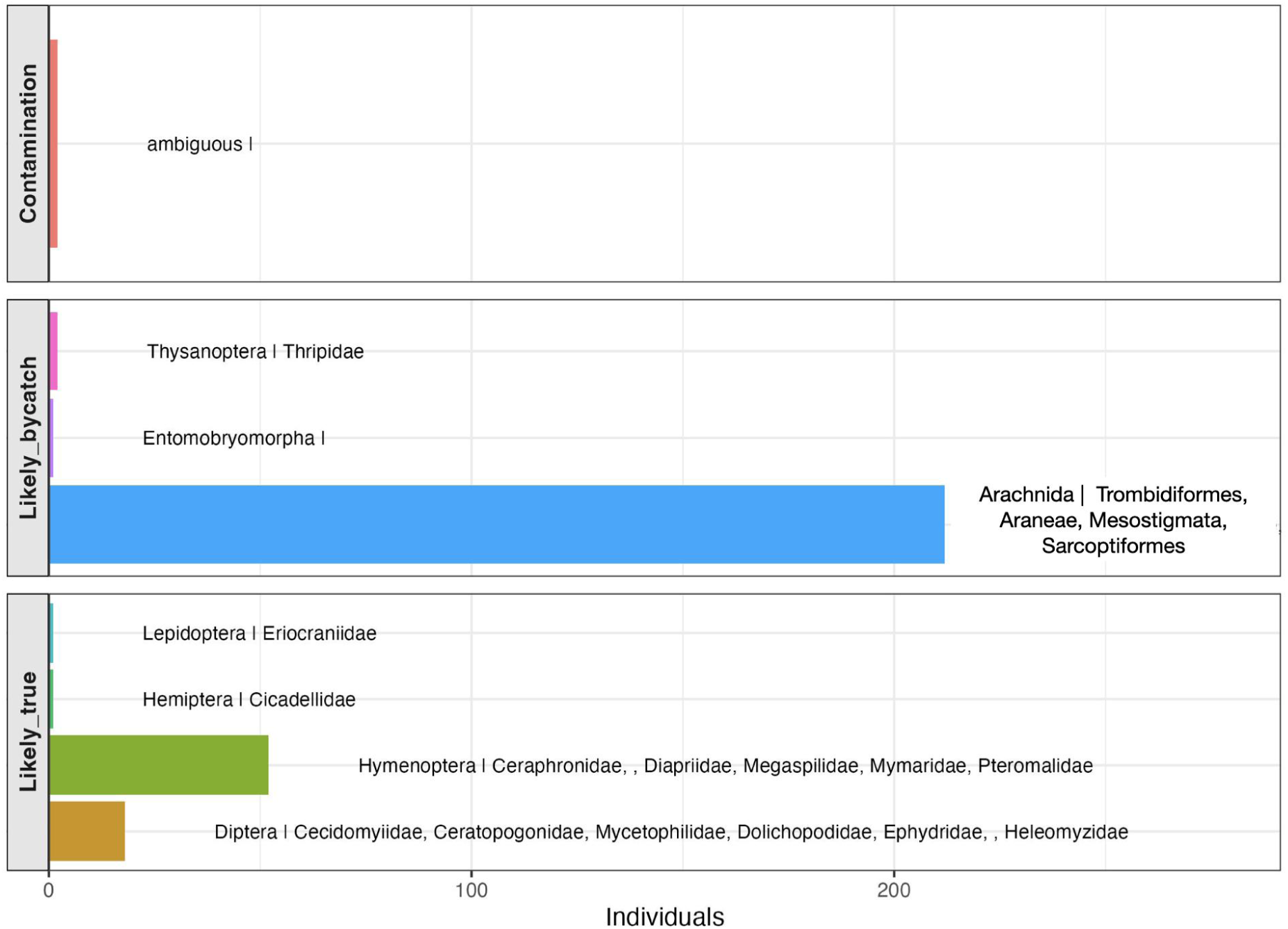
Taxonomic composition and classification of the 289 specimens that were successfully barcoded but did not match to metabarcoding dataset. The “Likely true” category consisted of (52 Hymenoptera: 17 haplotypes belonging to 13 clusters; 18 Diptera: 12 haplotypes belonging to 11 clusters, 1 Lepidoptera and 1 Hemiptera).

**Fig. S7.**
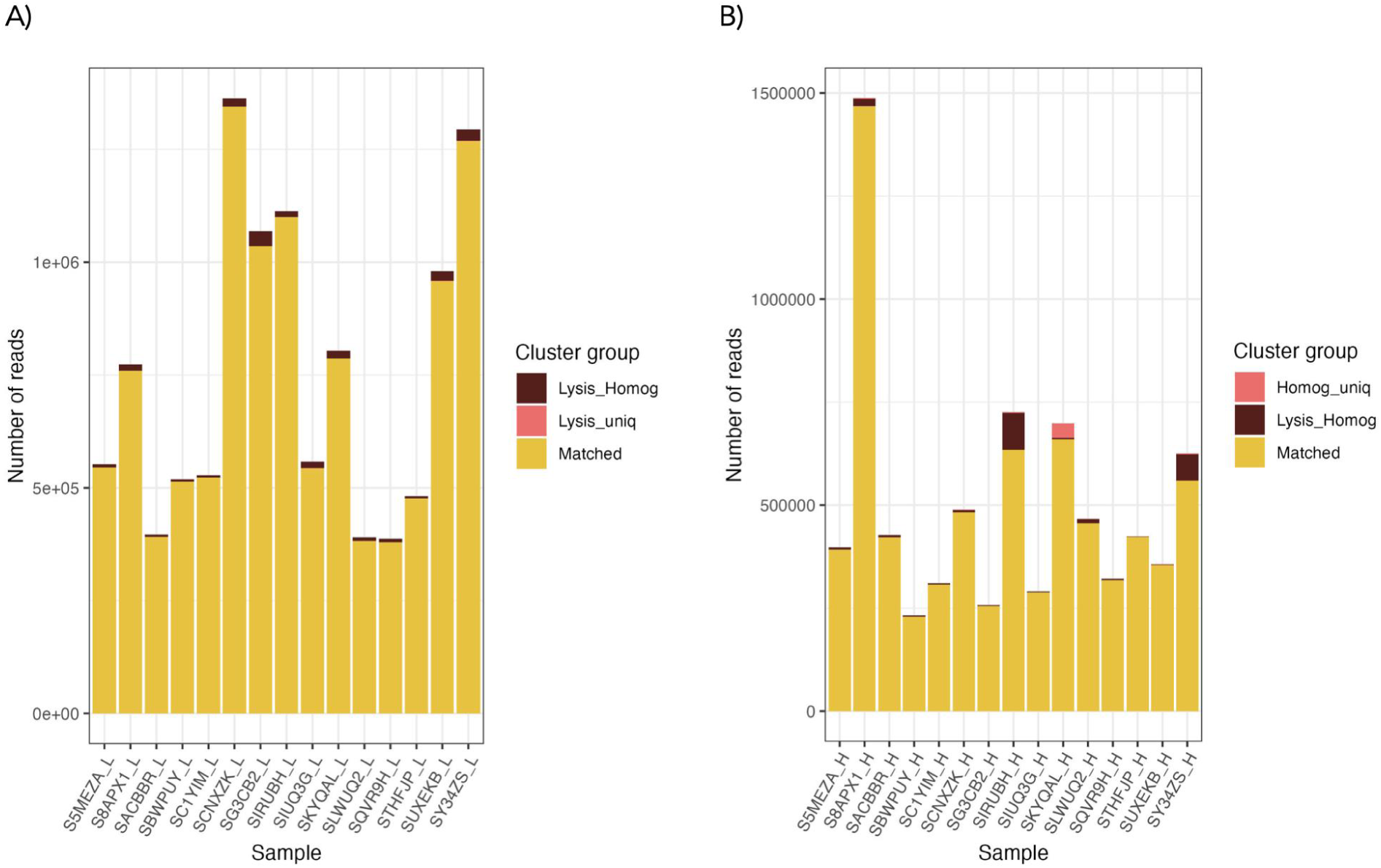
Histograms show the total number of metabarcoding read counts obtained per sample of the reference dataset after mild lysis (A) and homogenization (H). Yellow color indicates the proportion of read counts is accounted for by clusters that matched to the haplotypes of barcoded individuals. Brown color is the proportion of reads that represent clusters not matching to haplotypes and present in both metabarcoding datasets (mild lysis and homogenization). Bright pink shows the number of reads coming from clusters that were not matched to haplotypes and treatment-specific, i.e. found only in lysates (A), or only in homogenates (B).

**Fig. S8.**
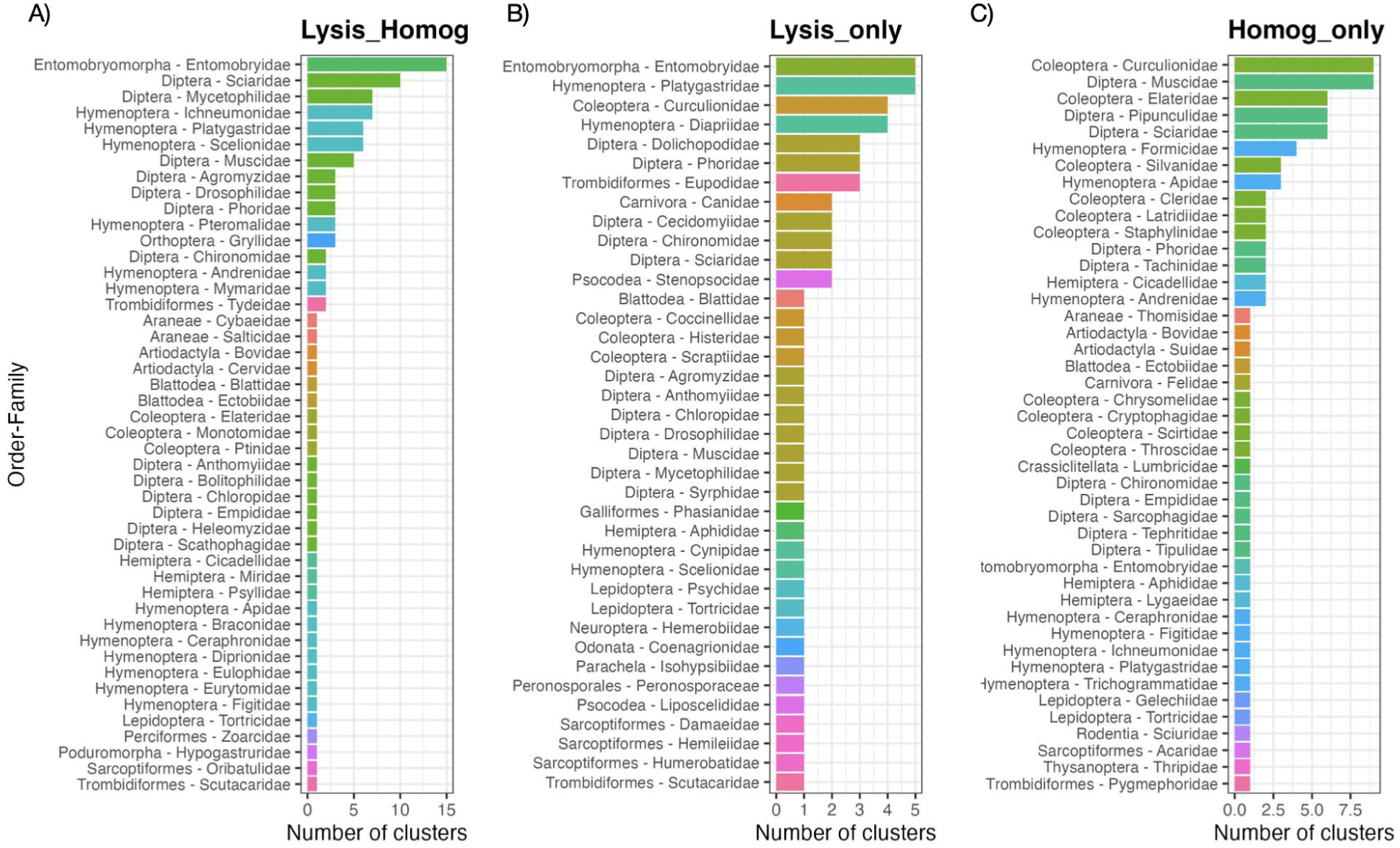
Summarizing taxonomic composition of clusters that were found in metabarcoding but not matched to individual barcoding results. A) Clusters found in both lysates and homogenates (N=109); B) Clusters found only in lysates (N=63); C) Clusters found only in homogenates (N=89).

**Fig S9.**
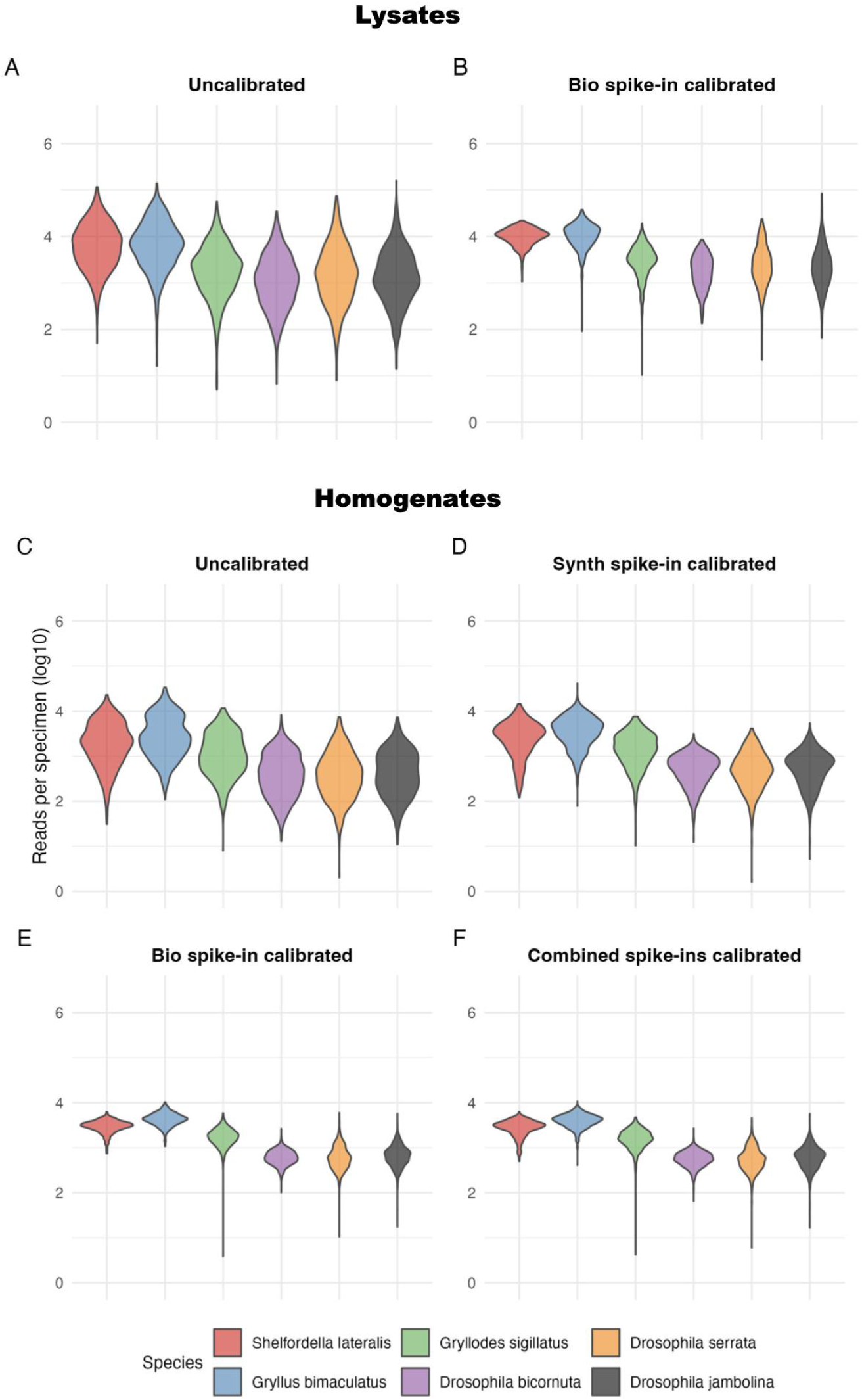
Calibration of read counts per specimen using biological or synthetic spike-ins to account for sample-specific effects. Data for 856 samples that underwent both lysis and homogenization. Log10-transformed read counts per specimen for six spike-in species across insect community samples. Panels (A and B) show lysate data, and panels (C–F) homogenate data (n=856). (A, C) Raw read counts. (B, D–F) Read counts after spike-in–based calibration. Panels (B, E) show calibration using biological spike-ins.

**Fig. S10.**
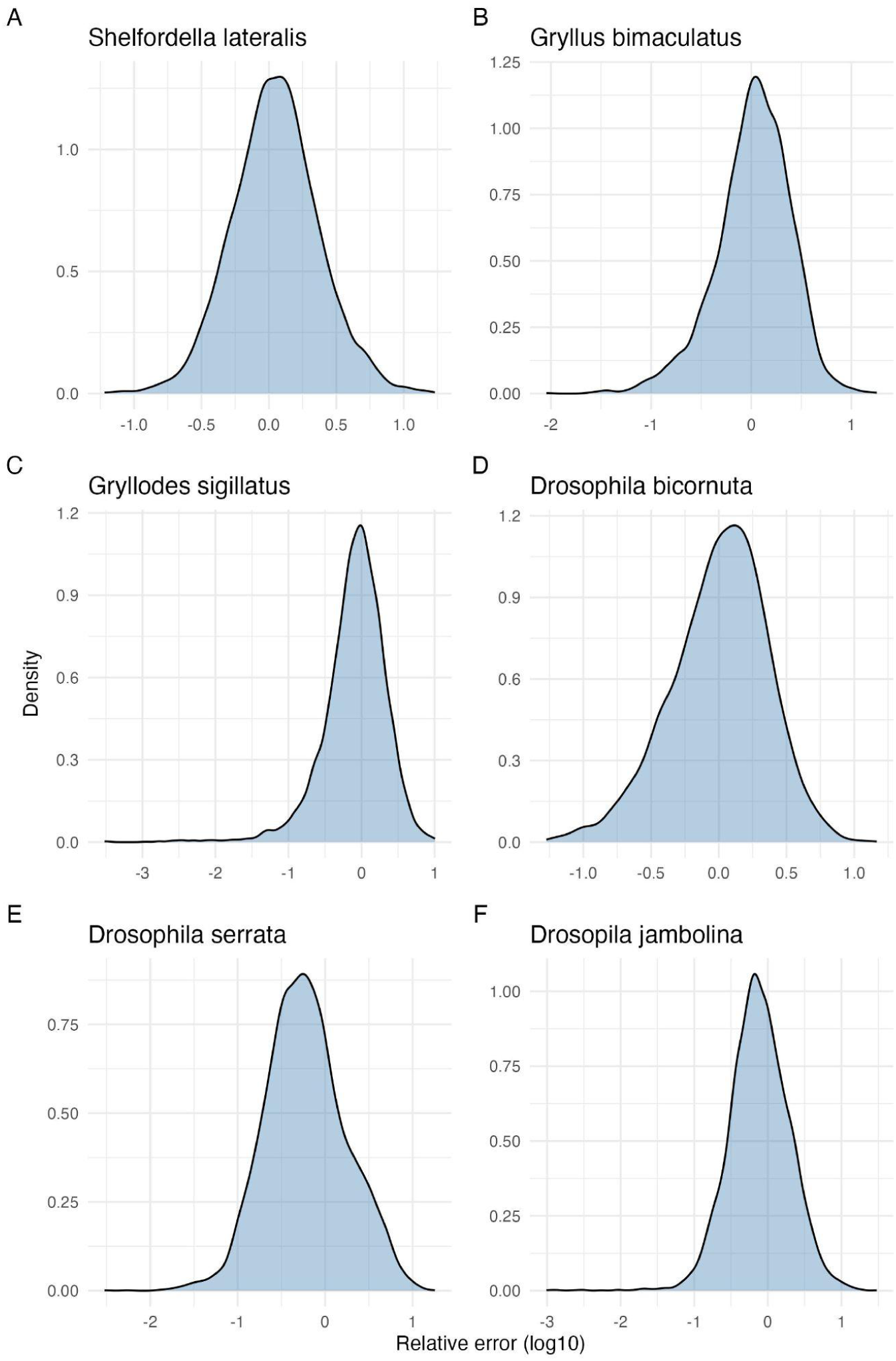
Kernel density plots of the log10 relative error for each biological spike-in species (panels A–F) in homogenate samples after leave-one-out bio-spike-in calibration. Values near zero indicate accurate recovery; the approximately symmetric shape indicates log-normally distributed residual error.

**Fig. S11.**
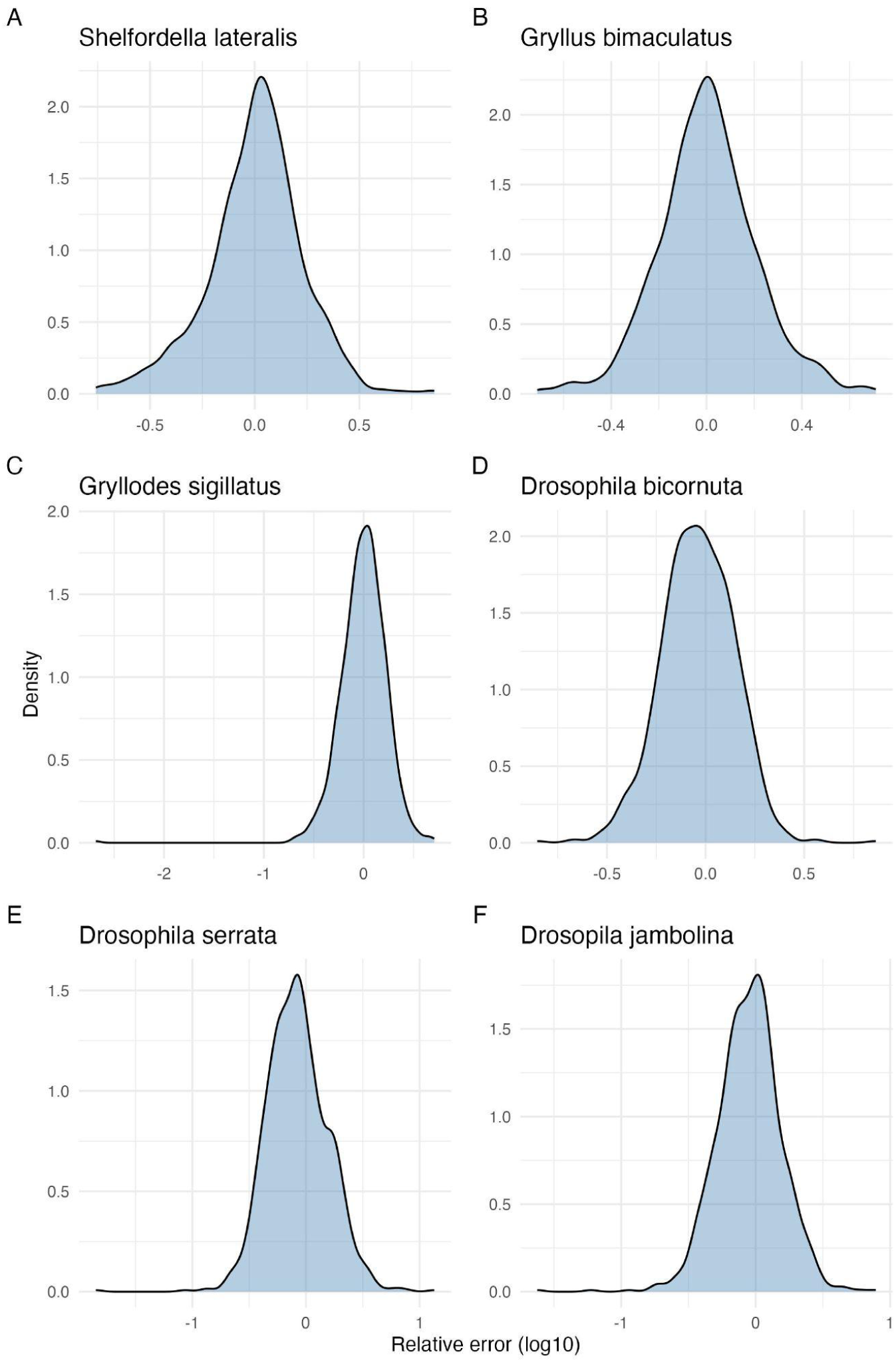
Kernel density plots of the log10 relative error for each biological spike-in species (panels A–F) in mild lysis samples after leave-one-out bio-spike-in calibration. Values near zero indicate accurate recovery; the approximately symmetric shape indicates log-normally distributed residual error.

**Fig. S12.**
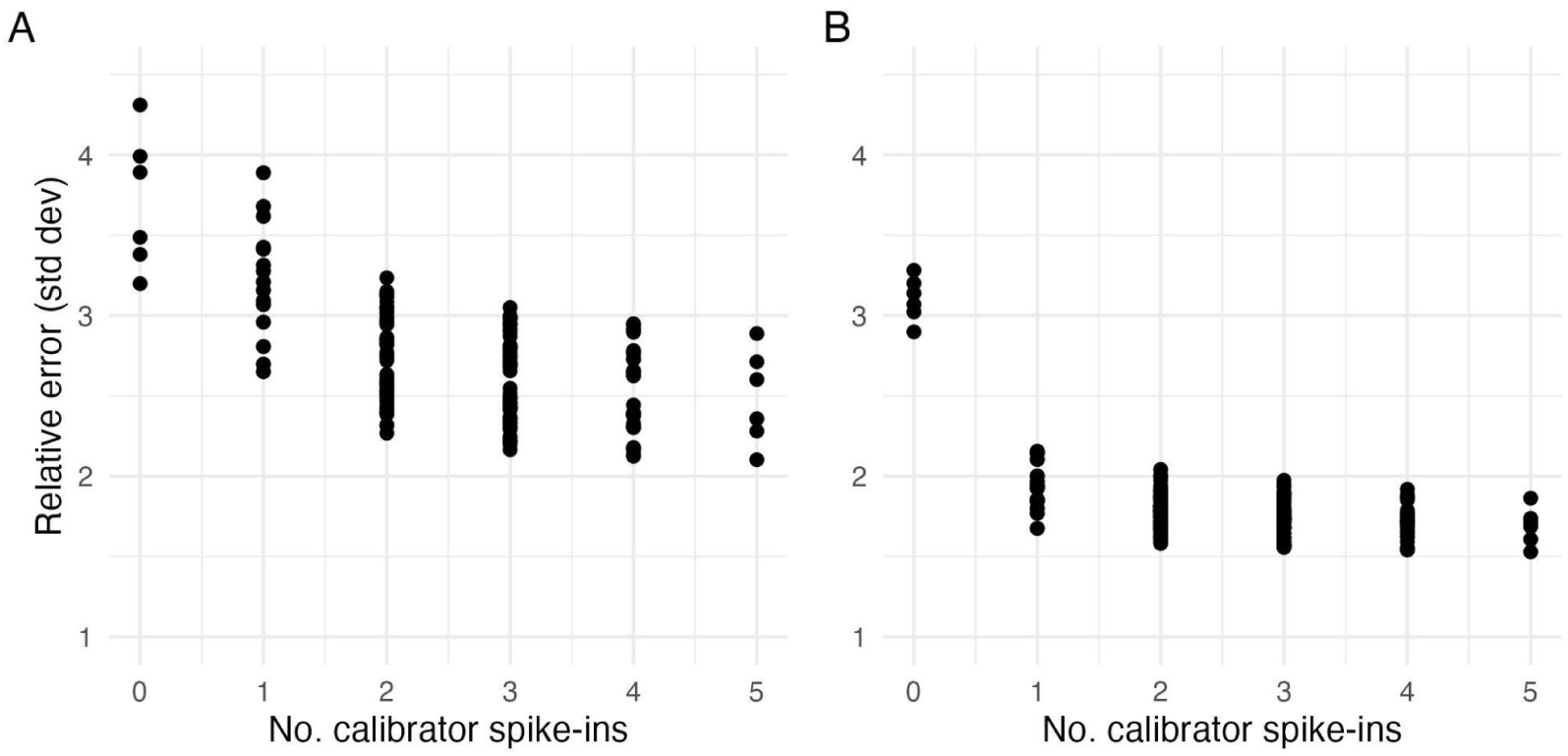
Effect of the number of calibrator spike-ins on relative error after mild lysis (A) and homogenisation (B). Each point shows the multiplicative standard deviation of the relative quantification error for one biological spike-in species evaluated using a specific subset of the remaining spike-ins as calibrators. The target species was always excluded from its own calibration.

**Fig. S13.**
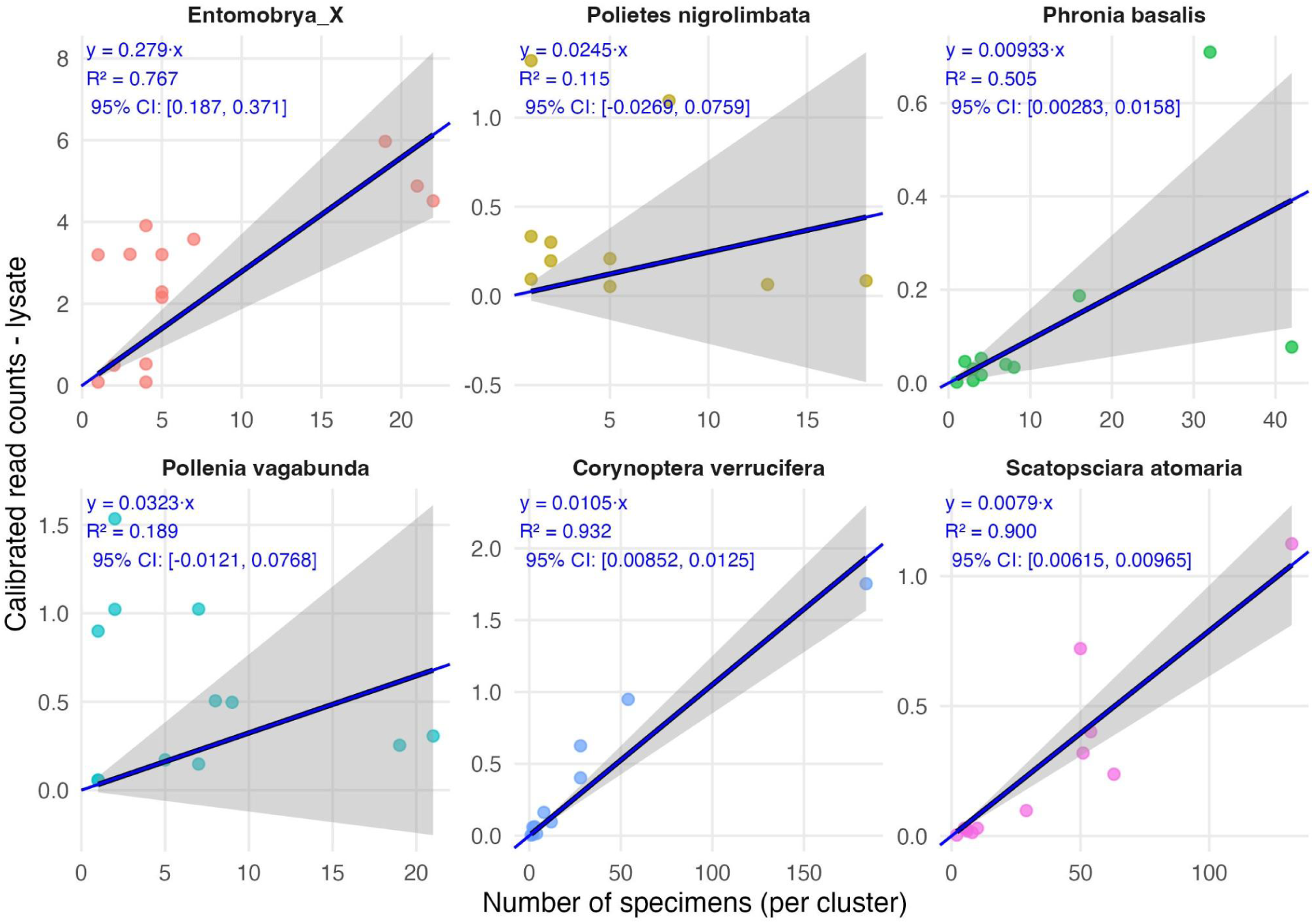
The plots above show, per cluster/species, the calibrated read counts from metabarcoding of the 15 reference samples over the number of specimens for that cluster for lysates. Only these six clusters in the 15 samples had both specimen count information (from barcoding) and non-zero read counts available in 10 or more overlapping samples.

**Fig. S14.**
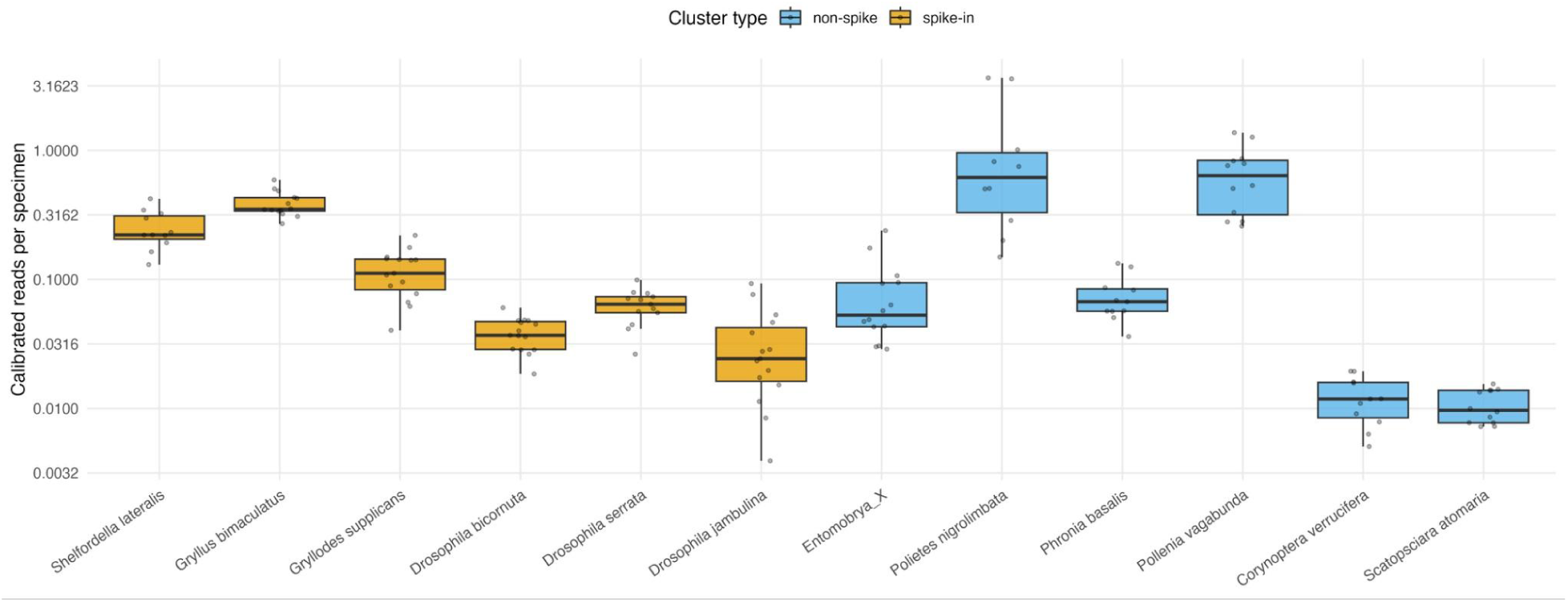
Boxplots showing the distribution of spike-in calibrated reads per specimens in the homogenate sample set. Orange boxplots show the biological spike-ins. Blue boxplots show non-spike-ins. Only taxa detected in at least ten samples were included, and only sample–taxon pairs for which both sequence reads and a positive ground-truth specimen count were available contributed to each box. Individual observations are shown as jittered grey points. The y-axis is on a log10 scale.

**Fig. S15.**
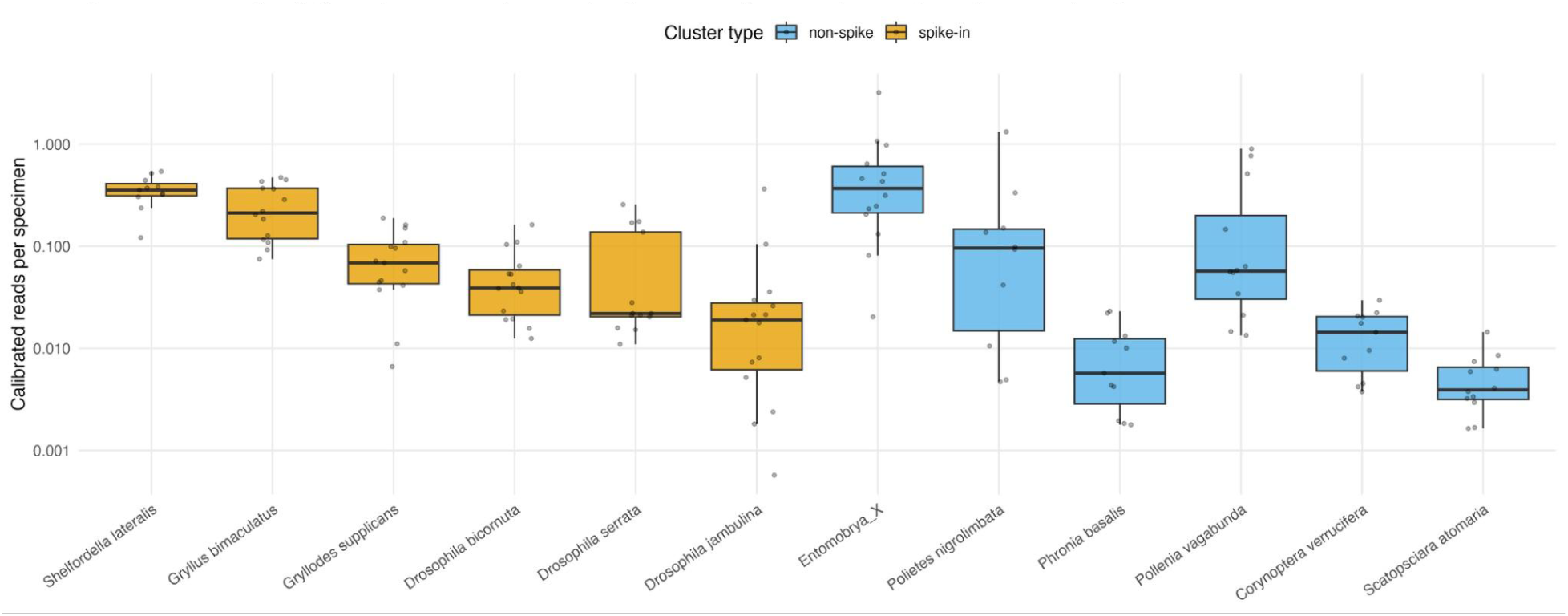
Boxplots showing the distribution of spike-in calibrated reads per specimens in the lysis sample set. Orange boxplots show the biological spike-ins. Blue boxplots show non-spike-ins. Only taxa detected in at least ten samples were included, and only sample–taxon pairs for which both sequence reads and a positive ground-truth specimen count were available contributed to each box. Individual observations are shown as jittered grey points. The y-axis is on a log10 scale.

**Fig. S16.**
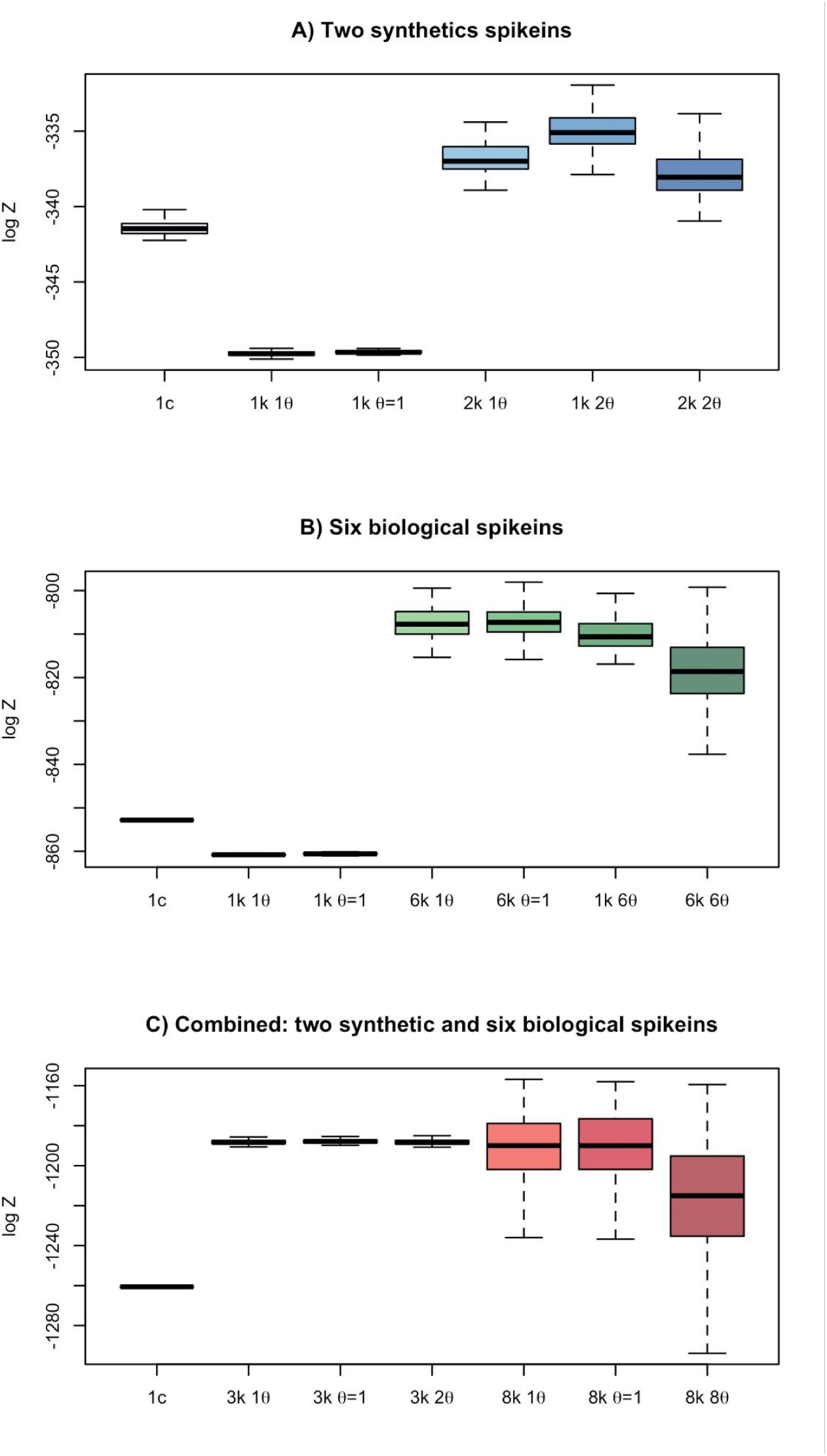
Normalization constants (log Z) for different abundance models, learning the distribution of the model parameters from the spikeins. There are three different sets of models: 1) Top panel in blue, using only the synthetic spikeins, 2) Middle panel in green, using only the biological spikeins, and 3) Bottom panel in orange, using a combined set-up of both synthetic and biological spikeins.

**Fig. S17.**
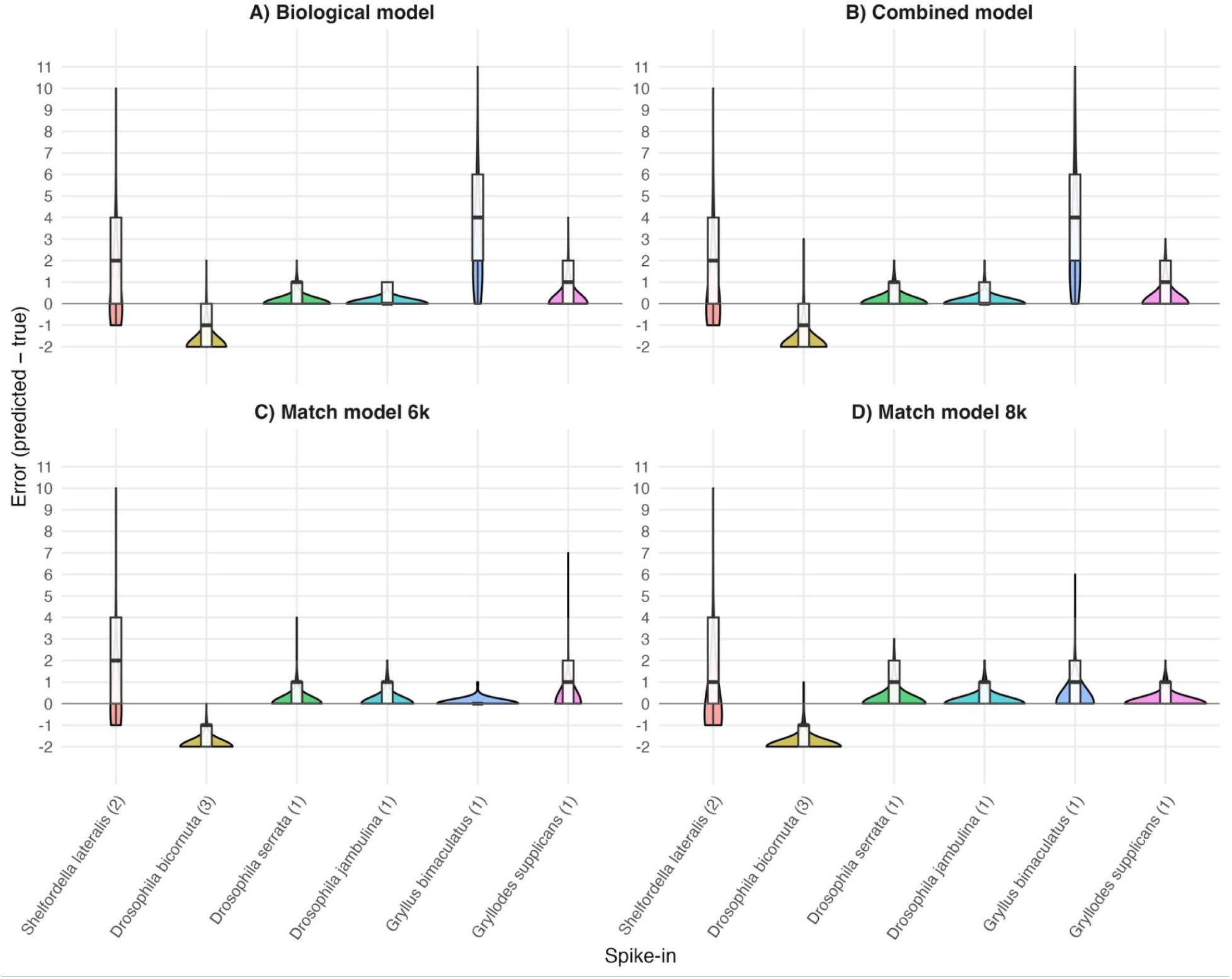
Error in the predictions of the specimen numbers of each of the six biological spike-in species. The upper panels show the biological model (**A**) and the combined model (**B**) using the same k value for all spike-in species used in the training. The lower panels show the biological model (**C**) and the combined model (**D**) using different k values for each of the spike-in species used in the training. Combined models used data from synthetic spike-in species in the training step.

**Fig. S18.**
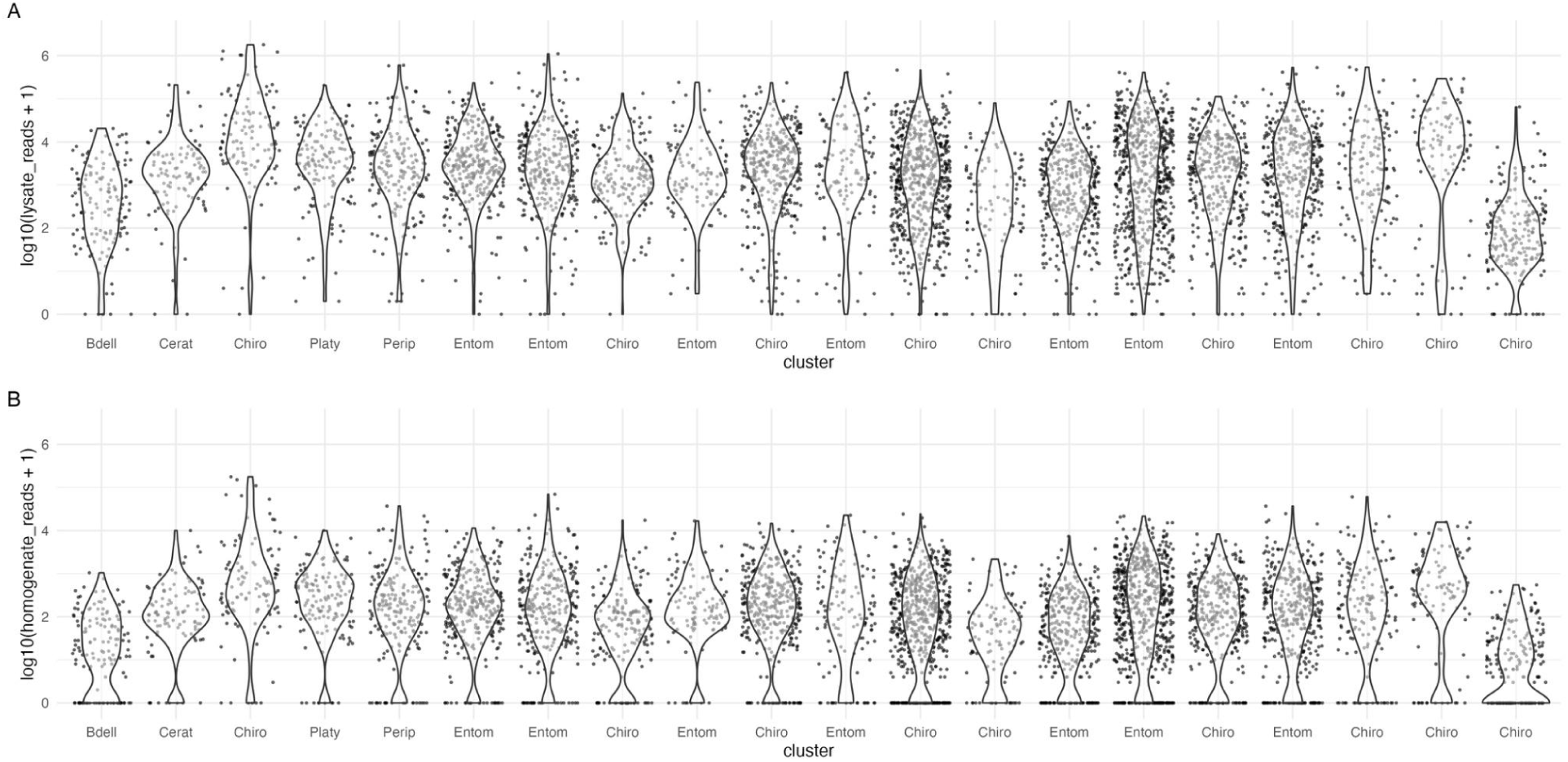
Top 20 clusters with largest surplus in lysates (top panel) when compared with homogenates (bottom panel).

**Fig. S19.**
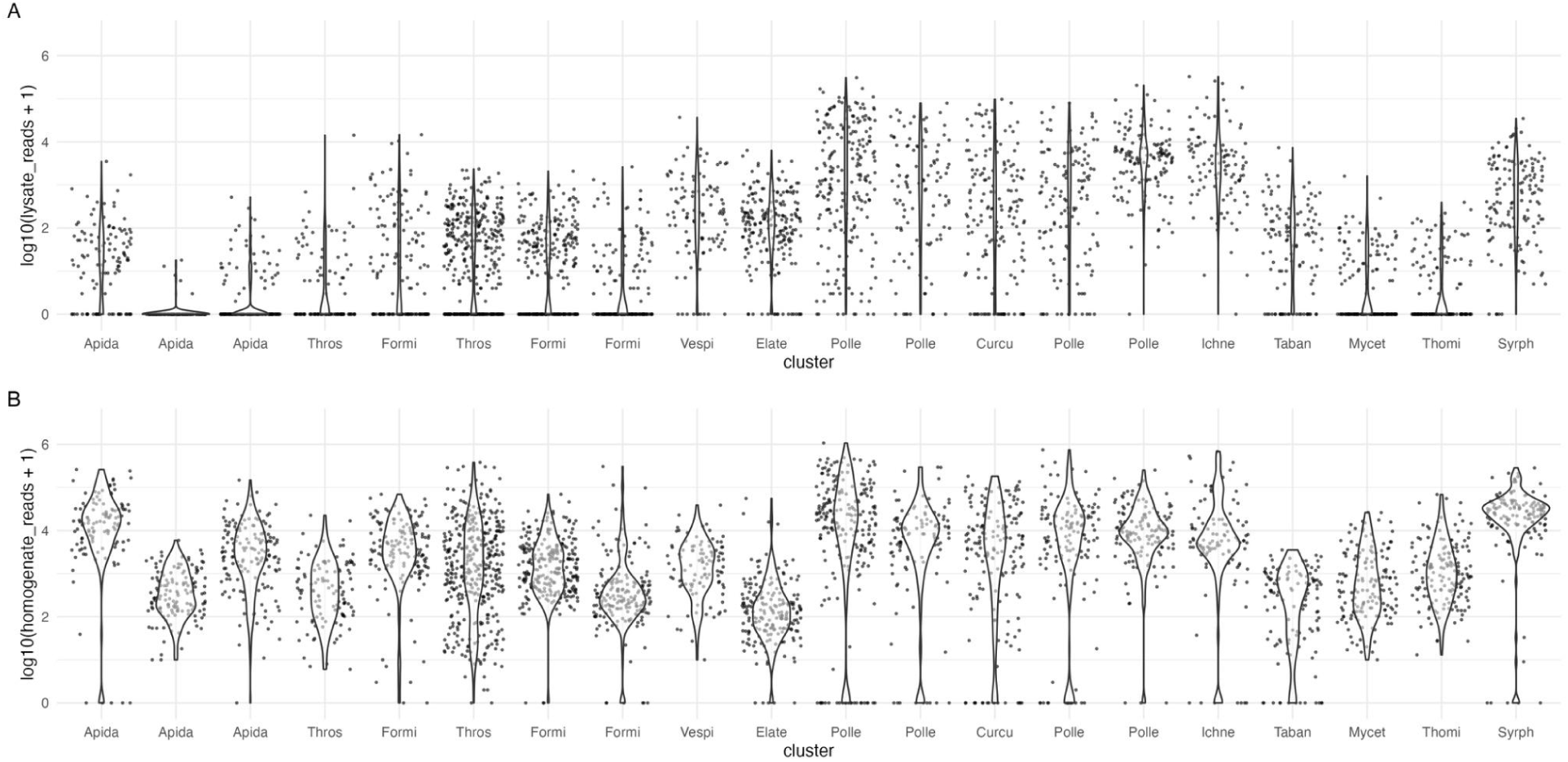
Top 20 clusters with largest surplus in homogenates (top panel) when compared with homogenates (bottom panel).

